# Structure of the human 80S ribosome at 1.9 Å resolution – the molecular role of chemical modifications and ions in RNA

**DOI:** 10.1101/2023.11.23.568509

**Authors:** Samuel Holvec, Charles Barchet, Antony Lechner, Léo Fréchin, S. Nimali T. De Silva, Isabelle Hazemann, Philippe Wolff, Ottilie von Loeffelholz, Bruno P. Klaholz

**Affiliations:** Centre for Integrative Biology (CBI), Department of Integrated Structural Biology, IGBMC (Institute of Genetics and of Molecular and Cellular Biology), 1 rue Laurent Fries, Illkirch, France; Centre National de la Recherche Scientifique (CNRS) UMR 7104, Illkirch, France; Institut National de la Santé et de la Recherche Médicale (Inserm) U964, Illkirch, France; Université de Strasbourg, Strasbourg, France; Architecture et Réactivité de l’ARN, CNRS UPR9002, Institute of Molecular and Cellular Biology (IBMC), Université de Strasbourg, 15 rue René Descartes, Strasbourg, France

## Abstract

The ribosomal RNA of the human protein synthesis machinery comprises numerous chemical modifications that are introduced during ribosome biogenesis. We present the 1.9 Å resolution cryo-EM structure of the 80S human ribosome resolving numerous new rRNA modifications and functionally important ions such as Zn^2+^, K^+^ and Mg^2+^ including their associated individual water molecules. 2’-O-methylation, pseudo-uridine and base modifications were confirmed by mass spectrometry resulting in a complete investigation of the > 230 sites many of which could not be addressed previously. They choreograph key interactions within the RNA and at the interface with proteins, including at the ribosomal subunit interfaces of the fully assembled 80S ribosome. Uridine isomerisation turns out to be a key mechanism for U-A base pair stabilisation in RNA in general. The structural environment of chemical modifications & ions is primordial for the RNA architecture of the mature human ribosome, hence providing a structural framework to address their role in healthy context and human diseases.

## Introduction

Chemical modifications of ribosomal RNA are introduced during biogenesis of the ribosome and have an important role in translation regulation with medical relevance. For the human cytosolic ribosome, the vast majority of chemical modifications comprises 2’-O-methyl (2’-O-Me) alterations of the ribose moieties and pseudo-uridines (Ψ)^1,2^. 2’-O-methylation is introduced by the methyltransferase fibrillarin guided by box C/D small nucleolar RNAs (snoRNA) to perform the enzymatic addition of a methyl group in a site-specific manner^1^, while isomerisation of uridines is catalysed by the pseudo-uridine synthase dyskerin (DKC1) within a complex of 4 core proteins guided by H/ACA snoRNA^2^. Such modifications have been characterized physicochemically or biochemically^3–7^, and more recently by RiboMethSeq^8,9^, AlkAnilineSeq^10^, HydraPsiSeq^11^, mass spectrometry^12^ etc. and listed in data bases such as Modomics^13^. In addition, other, yet less abundant modifications exist (∼5%) such as m^1^, m^6^ methylations etc. that occur on the nucleotide bases (introduced e.g. by the m^6^A methyltransferases METTL5^14^ or ZCCHC4^15^). There is growing evidence that such modifications can modulate ribosome function or cause antibiotic activity changes^16–27^. The hypothesis is that specific changes on “specialized” ribosomes may occur in a particular cellular context^28–31^, e.g. cell type or dysregulation. This has been observed recently for breast cancer in which certain 2’-O-Me are differently modified between tumors^32^, and similarly for pseudo-uridylation^33^, hence suggesting an important role for modifications in cancer^34,35^ in relation with the fact that the human ribosome represents a promising anti-cancer target^36,37^.Yet, the underlying molecular mechanisms and the role of modifications therein remain to be understood. Structural information in combination with biochemical characterisation of chemical modifications is essential to provide new clues into their role in various rRNA nucleotide positions.

The human 80S ribosome comprises the 60S subunit, which encompasses three rRNAs (28S, 5.8S and 5S), and the 40S ribosomal subunit that contains one single rRNA chain (18S). In an earlier study^38^ we performed a first visualization and annotation of chemical modifications in the human ribosome structure determined by cryo electron microscopy (cryo-EM)^39^ at an average resolution of ∼3 Å. While numerous modifications could be visualized, many more would be expected from data bases such as Modomics^13^ but could not be seen at the time, in particular 2’-O-Me and Ψ, the latter being limited in analysis by the possibility of resolving potentially associated water molecules; in addition, for some sites the chemical identity on nucleotide bases was unclear and remained to be addressed. Also, according to electrostatic potential map calculations^40^, some previously non-annotated sites on the 28S rRNA at the N7, O6 or O4 atoms of some guanines (G), adenines (A) or uridines (U), respectively, appeared more likely to instead represent Mg^2+^ ion positions but this remained to be confirmed experimentally; positive charges tend to generate larger densities in the cryo-EM map and may sometimes erroneously suggest the presence of a chemical bond. Better resolved maps would help overcoming such limitations. More generally, identifying the position of typically RNA-associated ions such as Mg^2+^ and K^+^ and localizing water molecules in the structure would be important functionally. Using the latest-technology cryo electron microscope setups (Titan Krios G4 from Thermofisher Scientific and CRYO ARM 300 from Jeol, see methods), including latest-generation energy filters and direct electron detection cameras, combined with advanced image processing, sorting and extensive 2D & 3D focused classification & refinement procedures^41–50^ to push resolution even further including with higher order aberration correction, we now succeeded in breaking the 2 Å resolution barrier for the structural analysis of the 80S human ribosome, a non-symmetrical, rather flexible complex^36,38,51,52^ that compared to the bacterial ribosome comprises long disordered rRNA expansion segments and lacks stabilization when binding antibiotics. The structure provides unprecedented detailed insights into the overall RNA and protein structure of the human ribosome, the localisation of ions, ligands and water molecules and in particular chemical modifications of the rRNA in both 40S and 60S ribosomal subunits. In addition, to corroborate the identification of sites we used a mass spectrometry (MS) based method developed for the characterisation of RNA modifications^52^, which was adapted and extended to the analysis of rRNA modifications^54,55^. To clarify sequence inconsistencies in data banks we performed sequencing of the 28S, 18S, 5.8S and 5S of the rRNAs to unify the annotation of nucleotides and chemical modifications. Together, this allowed analysing the precise structural environment and function of chemical modification in the human 80S ribosome structure.

## Results

### High-resolution structure determination of the human ribosome

Human ribosomes were isolated from HeLa cells and flash-frozen on cryo-EM grids from which those with thin amorphous ice were selected for data collection using cryo electron microscopes equipped with energy filters and direct electron detection cameras (see methods & **Extended Data Fig. 1**). Initial image processing including particle sorting and 3D classifications provided a first map at 2.3 Å resolution. A particular attention was then given towards refinement of parameters such as contrast transfer function (CTF) and higher-order aberrations using optic groups followed by particle polishing (see methods) to account for minor imaging abnormalities and particle movements, reaching 2.1 Å resolution (**Extended Data Fig. 2**). Further focused and multi-body refinements (**Fig. 1a**) yielded 1.9 Å average resolution, with an average resolution of 1.9, 2.0 and 2.1 Å for the 60S ribosomal subunit and the body and head regions of the 40S ribosomal subunit, respectively. An additional final refinement on each body using partial signal subtraction further improved features in the cryo-EM map (**Extended Data Figs. 2 & 3**; see methods). Local resolution analysis shows that many regions extend to 1.7-1.8 Å resolution (**Fig. 1a**).

**Figure 1.**
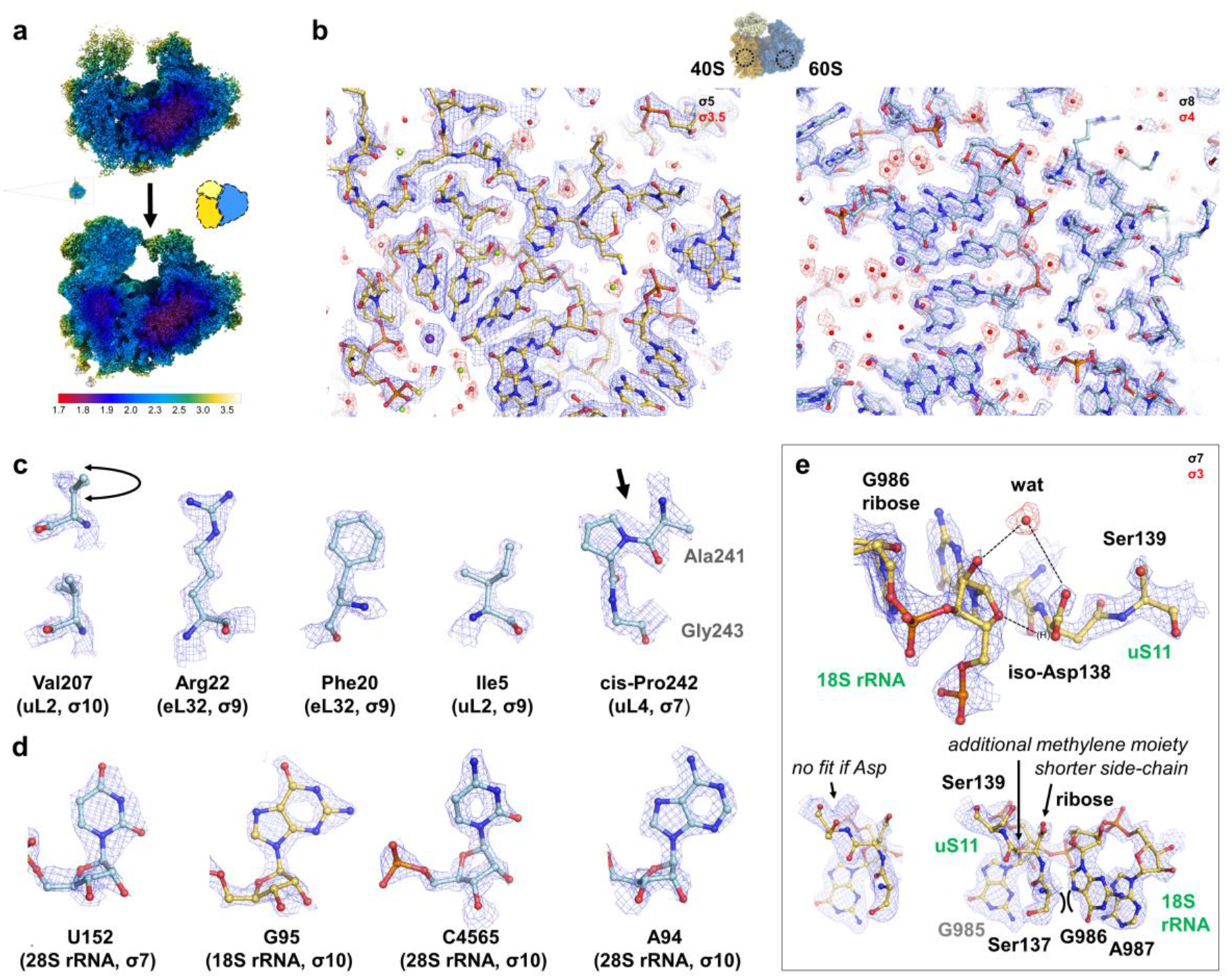
High-resolution features, ions and water molecules in the human 80S ribosome. **a** Local resolution estimated with the Relion software before (top) and after (bottom) multi-body refinement (before final partial signal subtraction, see methods). The different masks used for the multi-body refinement are shown with dotted-lines. **b** Global views of the 40S and the 60S regions illustrating the quality of the map (sigma level indicated in top right corners). **c** Examples of several amino acids from ribosomal proteins uL2, eL32 and uL4. The resolution gives insight on the exact positioning of the di-methyl moiety of a valine side-chain (shown before and after flip and fitting), allows to see the zig-zag of an Arg side chain, allows to see the hole in the aromatic side chain of a Phe, gives the possibility to fit the ethyl and methyl moieties of an Ile and to distinguish a cis-Pro from a trans-Pro. **d** Examples of nucleotides in the 28S and the 18S mRNA. The high-resolution features resolve the ring structure of purines and pyrimidines side-chains and in riboses. **e** Example of a modified iso-Asp138 in ribosomal protein uS11 in the 40S region. An additional methylene moiety is visible in the backbone and the side-chain is shorter. The iso-Asp side chain interacts with the G986 ribose through H-bond to stabilize the local region. In the bottom left corner, the Asp before modification is shown, making it impossible to fit in the cryo-EM map.

We then built an atomic model based on our previous lower resolution structure^38^ (PDB ID 6QZP), but now refined in much more detail to account for precise side-chain conformations on ribosomal proteins and rRNA thanks to the high-resolution features of the cryo-EM map (both for the 40S and 60S subunits, **Fig. 1b-d**; see methods). The map resolves numerous chemical modifications, ions and water molecules (annotated as described in the following), which were built and included in the atomic model and refined using the Phenix software^56^ to obtain good geometric refinement parameters (**Extended Data Fig. 1**).

### Specific features, post-translational modifications and ions in the high-resolution structure of the human ribosome

The structure provides unprecedented insights into many fine details. For instance (**Fig. 1c**), valine side-chain up and down conformations can be differentiated, side-chain conformations of isoleucine residues can be resolved because the orientation of methyl and ethyl moieties can be distinguished, and cis-prolines can be identified. High-resolution features can be observed on the ring structure of nucleotide bases and ribose moieties of the rRNA backbone (**Fig. 1d**). Chemical modifications can be found on numerous rRNA residues (as discussed later; **Extended Data Figs. 4 & 5**) but few are visible on the ribosomal proteins (this does not exclude the possibility that more exist) including methyl-arginine (ribosomal protein eS19), hydroxy-histidine (uL15, important for translational activity^57^) and methyl-histidine (uL3) residues (**Extended Data Fig. 6**), consistent with post-translational modifications reported in the Uniprot database. A particularly interesting example is iso-aspartate found in the universally conserved ribosomal protein uS11 (**Fig. 1e**), a residue that contains an extra methylene moiety in the backbone and thereby modifies the local geometry; this modification has been observed in the *E. coli* ribosome^58^ and in plants^59^ (tomato; also visible in tobacco^60^ but not modelled), hence confirming the conservation of this rather peculiar post-translational modification across species including human. The shorter methylcarboxylate side-chain of iso-Asp138 packs smoothly against the ribose moiety of G986 and forms a water-mediated hydrogen bond), which would not be possible with a standard aspartate residue (**Fig. 1e**). The functional meaning of this modification remains to be understood (for other proteins, it is linked to diseases^61,62^), but structurally it stabilizes the interaction between the 135-140 extended loop region of uS11 and the single-stranded 986-989 region of the 18S rRNA on the 40S platform next to the tRNA exit (E) site and the 5’ end of the mRNA channel.

Many additional small densities can be found in direct vicinity of nucleic acid bases, phosphate and ribose moieties and amino acids which potentially correspond to ions and water molecules (**Fig. 2**), and some polyamines (2 endogenous spermidine molecules as no polyamines were added in the buffer; **Extended Data Fig. 7**). Due to the complexity of the structure and the sheer amount of individually resolved densities we first placed ions and then searched for water molecules (see methods). To annotate ions in a reliable manner we considered both the strength of the map density and their characteristic coordination known from chemistry: tetrahedral for Zn^2+^ ions (typically in a surrounding environment of Cys residues), octahedral for Mg^2+^ ions, and higher coordination (e.g. 8-12) for K^+^ ions (the latter show a stronger density than other ions; detailed below & see methods), hence considering distances and angular geometry for each ion type in its environment. Taken together, 9 Zn^2+^, 294 octahedral Mg^2+^ and 107 K^+^ ions could be annotated (**Fig. 2a**), along with ∼16000 water molecules (using an automated procedure that combines density search and coordination with hydrophilic residues considering potential hydrogen bonds using SWIM^63^ and Phenix^56,64^ software; see methods).

**Figure 2.**
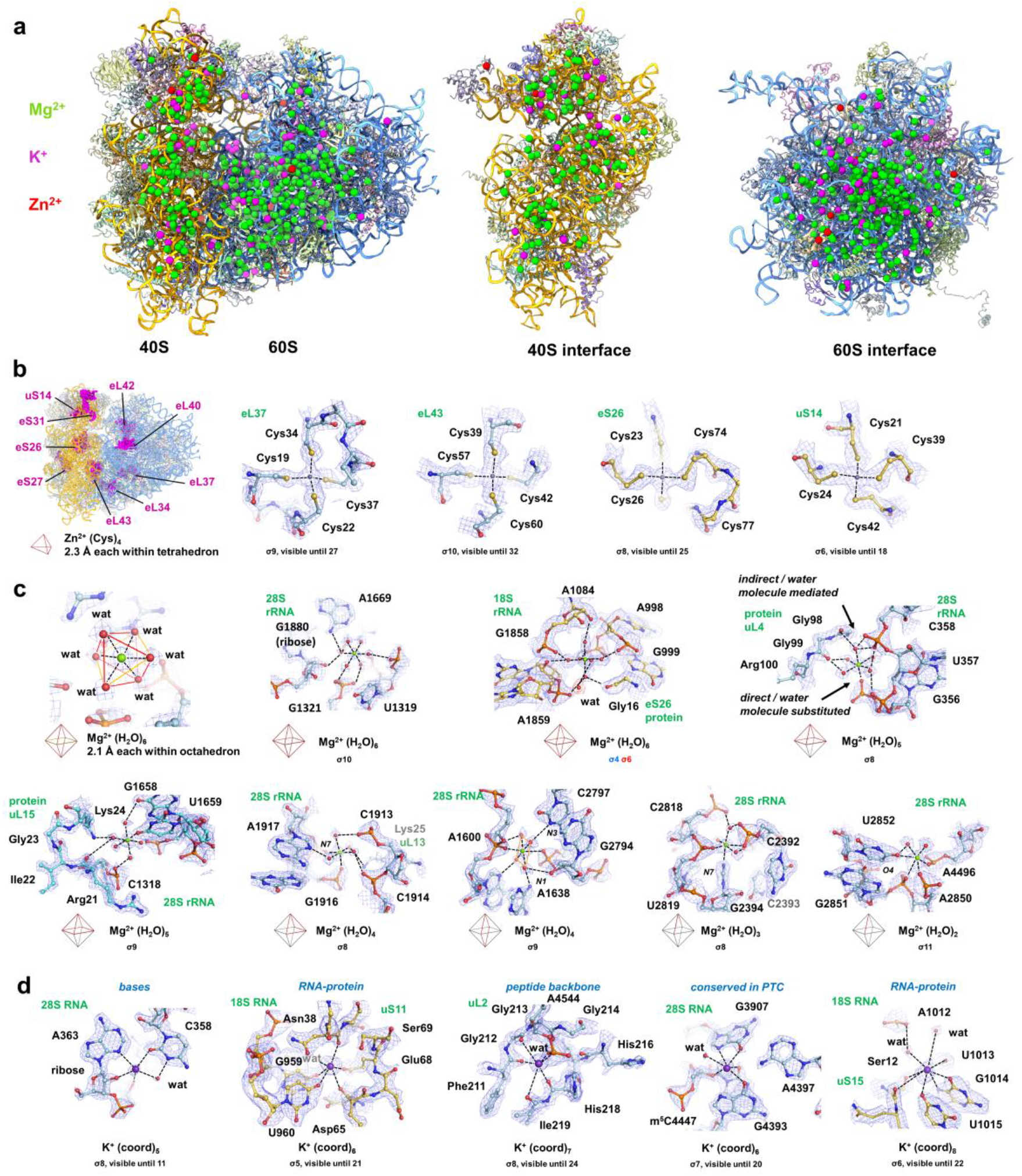
Assignment of ion coordinations in the human 80S ribosome. **a** Overall structure of the human ribosome with highlighted positions of Mg^2+^, K^+^ and Zn^2+^ ions. **b** (*Left*) Overall positioning of the nine Zn^2+^ binding ribosomal proteins and (*right*) four examples of the Zn^2+^ binding sites in the 60S large ribosomal subunit and the 40S small ribosomal subunit. Respective s-levels used to prepare figures and s-levels at which the ions are still detectable are indicated. **c** Selection of sites with octahedral water-coordinated Mg^2+^ ions. The number of coordinated water molecules can range from six to two depending on the number of direct interactions of the Mg^2+^ ions with oxygen atoms of ribosomal RNA or proteins. Mg^2+^ can coordinate ribosomal components directly or via a water molecule. Respective s-levels used to prepare figures are indicated. **d** Selection of sites that show coordinated K^+^ ions. K^+^ was found to have up to eight coordination partners, which include ribosomal proteins, water molecules or rRNA. The conserved K^+^ ion in the PTC identified previously^68^ was also detected in the present structure. Respective s-levels used to prepare figures and s-levels at which the ions are still detectable are indicated.

Coordinated Zn^2+^ ions can be seen in several ribosomal proteins (eL34, eL37, eL40, eL42, eL43, uS14, eS26, eS31). Several are very well resolved and show the characteristic tetrahedral configuration by 4 cysteine residues with a typical short distance of 2.3 Å between the cysteine sulphur atoms and the Zn^2+^ ion (**Fig. 2b**). Ribosomal protein eS27 exhibits a cysteine-preconfigured tetrahedral site (but no Zn^2+^ density is visible there, suggesting a weak binding site). With the exception of uS14, the Zn^2+^-binding sites are found primarily in eukaryote-specific ribosomal proteins. This suggests that the evolutionary addition of proteins in eukaryotes (as compared to bacterial ribosomes) required additional stabilization of the protein fold by a coordinating Zn^2+^ ion, possibly related with the fact that all the Zn-binding eukaryote-specific ribosomal proteins are surface-exposed (**Fig. 2b**) which may entail additional stabilization.

Mg^2+^ ions are readily identifiable thanks to their characteristic octahedral coordination with water molecules in 2.1 A distance from the central ion (**Fig. 2c**). This is possible because the associated individual water molecules can often be resolved in the map or the octahedral hydration shell becomes visible at lower contour level when a given site is less stable (as illustrated by variable temperature factors). Hexahydrated Mg^2+^ ions were fitted as octahedrons into the cryo-EM map taking into account the possible substitution of one or more water molecules by a polar substituent (refinement of the octahedrons was restrained in Phenix, see methods). This leads to variable configurations within the octahedrons, either with 6 water molecules or variations with an increasing number of residues (e.g. 1, 2, 3 or 4) taking every time the place of one octahedron corner and leaving respectively 5, 4, 3 or 2 water molecules (**Fig. 2c**). Substituents are found to be exclusively oxygen atoms, e.g. from negatively charged phosphate groups (which favours charge compensation), 2’-hydroxyl groups of ribose moieties and oxygen atoms of nucleotide bases and amino acids (**Fig. 2c**). The analysis confirms our previous ESP-annotated Mg^2+^ sites^40^ that can now be resolved as such at N7, O6 and O4 positions of some G, A and U in the 28S rRNA (**Extended Data Fig. 8**) instead of an initially assumed chemical modification^38^; these nucleotide base positions function as weak Lewis bases and therefore provide only partial charge compensation as compared to a phosphate group. Importantly, the possibility to resolve the associated water molecules throughout the structure reveals that in many cases the Mg^2+^ ion contacts to the neighbouring residues are water molecule mediated, *i.e.* even phosphate groups often interact with the Mg^2+^ ion indirectly via a water molecule rather than directly as might be assumed classically (**Fig. 2c**). These Mg^2+^ ions generate interaction links between different structural elements (e.g. nucleotides G1880-A1669-U1319, A998-A1084-A1859, A1600-C2797 and U2852-A4496; 28S rRNA; **Fig. 2c**) that are relatively far away in sequence and in distance (∼4-8 Å, i.e. much beyond hydrogen bonding distance), hence stabilizing the rRNA fold.

The characteristic higher-order coordination of K^+^ ions is visible as a recognizable feature (notably stronger density than a water molecule or other ions because of its positive charge, no close coordination by 6 regular water molecules in contrast to octahedral Mg^2+^, which at lower resolution - where water molecules cannot be seen may lead to falsely assigned K^+^ ions) together with the more variable bonding distance and irregular geometry resembling a square anti-prismatic (**Fig. 2d**) as seen in potassium channels and in the *T. thermophilus* ribosome^65,66^. K^+^ ions are typically found at interface areas between rRNA bases and amino acid side-chains (**Fig. 2d**), or involving 2’-OH ribose moieties (e.g. A363 in 28S rRNA, **Fig. 2d**), or stabilizing protein loops (Gly213 region of ribosomal protein uL2 which comprises 3 consecutive glycine residues; **Fig. 2d**). Their environment almost exclusively comprises oxygen atoms representing weaker Lewis bases (mostly amino acid backbone and nucleic acid keto groups and few hydroxyl groups, i.e. usually not phosphate groups). The role of K+ ions in stabilizing these structural elements -typically intrinsically less stable single-stranded rRNA or protein loop regions - implies also an important role in translation, which is consistent with the observation that ribosomes lacking K^+^ ions show reduced poly-Phe synthesis and increased thermal instability^67^. In this context, there is a characteristic K^+^ binding site located at the PTC, which bridges G3907, G4393 and m^5^C4447 of 3 different 28S rRNA segments (**Fig. 2d**). The interaction with the phosphate moiety of m^5^C4447 creates a charge compensation, while coordinating with the 6-keto group of G4393 and the 6-keto and N7 positions of G3907. Interestingly, this K^+^ position is also observed in the bacterial ribosome structure^68^, hence suggesting a conserved role across species. Furthermore, we could also identify K^+^ binding to the N7 position of G1484 showing some exchangeability between Mg^2+^ and K^+^ at this base moiety.

## MS, rRNA sequencing & structural analysis of chemical modifications

RNA modifications are chemical changes that usually result in a mass shift, hence MS allowed to detect and determine the modifications placement on the RNA sequence. MS analysis of chemical modifications was done on the 18S, 28S, 5.8S and 5S rRNAs using site-directed synthesized oligos combined with enzymatic cleavage with four different RNases (T_1_, A, U_2_ and MC1) and high-resolution LC-MS/MS analysis of RNA fragments allowed to determine the modified positions on the RNA sequence (see methods; **Extended Data Figs. 9-11**). 2’-O-Me sites were identified by MS and cross-validated with the cryo-EM structure, in particular for the sites not previously observed (e.g. Um428 from 18S or Um4498 from 28S rRNA). Ψ sites were identified by MS using specific acrylonitrile labelling to generate a corresponding mass shift; identified sites were confirmed in the structure based on the characteristic interaction of water molecules or rRNA phosphate groups of the rRNA backbone with the exposed N1 position of Ψ’s. Sites not found by MS were annotated according to previous identifications^9,11,12,13,31,69,70,71,72^ and cross-checked in the structure.

To overcome confusing differences in rRNA sequence numbering as found in data banks we performed rRNA sequencing (based on reverse-transcription followed by PCR, see methods); this includes a sequence correction from G4910 to adenine in the 28S rRNA, which is visible also in the structure (**Extended Data Fig. 12**) hence providing a unique consistency validation between sequence, structure and MS analysis of the rRNA using the same sample. All chemical modifications were annotated on the rRNA sequences (**Extended Data Figs. 13 & 14**). This now allows an unambiguous annotation of rRNA residues both in the structure and with regards to chemical modifications found in the literature. This turns out to be greatly helpful to unify the entire annotation of modifications sites on the basis of a unique numbering (99 2’-O-Me sites, 109 Ψ, and 12 base modifications such as m^1^acp^3^, m^1^, m^3^ etc. for the structural data and respectively 6 and 9 seen only from the MS data; total of 235 sites including 105 2’-O-Me & 118 Ψ and 12 base modifications). The MS-identified 2’-O-Me sites and Ψ sites are overall consistent with previous identifications^9,11,12,13,31,69,70,71,72^ (**Suppl. Data File 1**). New sites found by MS and also observed in the structure include for 18S rRNA Ψ63, Ψ100, Ψ300, Ψ1003, Ψ1360, Ψ1596. Those newly found by MS but not observable in the structure due to weakly resolved regions include for 18S rRNA Ψ366, Ψ514, Ψ556, Ψ667, Ψ769/770 (the only one on an rRNA expansion segment), Ψ804, Ψ1061, Ψ1186, and Gm1946 and Ψ5039 for 28S rRNA; Ψ897 listed in Modomics^13^ is not found by MS and is not visible in the structure; Ψ1136 found in Modomics is not found by MS but is visible in the structure. Together, the combined high-resolution structural analysis, sequencing and MS-based validation allows a full annotation of chemical modification sites.

The structure comprises in the 28S, 18S and 5.8S rRNA’s, respectively, 62, 37 and 2 2’-O-Me and 58, 49 and 2 Ψ modifications, most of which are confirmed by our MS analysis and otherwise identified by previous studies^9,11,12,13,31,69,70,71,72^. 5.8S rRNA carries 4 modifications all of which were found by MS, while none were found in the 5S rRNA (nor predicted nor visible in the structure). Compared to the full set of modifications found by MS and/or previous studies, 12 2’-O-Me and 13 Ψ modifications are not visible in the structure (e.g. when less resolved in the periphery of the complex). Interestingly, the MS data of the modification m^1^acp^3^ψ1248 (18S rRNA) reveal an extra mass corresponding to the addition of a cyanoethyl group. Considering that the only reactive nitrogen atom available is located on the modification, this indicates the formation of a cyanoethylated acp3 adduct due to the chemical treatment, which to our knowledge has not been observed before (see methods).

### Ribose 2’-O-methyl modifications in the human ribosome

2’-O-methylations are found fairly widespread throughout the structure, with rather many in the 40S compared to the 60S ribosomal subunit, including clusters at functional centres such as the peptidyl transferase centre (PTC) and the decoding centre (DC), but only few on the solvent side of the 60S ribosomal subunit (**Fig. 3**). The 5.8S rRNA has two 2’-O-methylation sites as confirmed by MS (**Fig. 3b & c**), both of which do not show particular contacts to neighbouring residues alike some sites in 18S and 28S rRNA; this may imply that other cellular components interact there, in particular when the sites are solvent-accessible as is the case for the 5.8S rRNA sites. However, many others do show contacts with neighbouring residues, which translate into van der Waals contacts due to the hydrophobic nature of a methyl group. Hydrophobic contacts typically involve hydrophobic side-chains of amino acids (**Fig. 3g, h, i, l**) or the aromatic face of nucleotide bases (**Fig. 3e, k**) or between several 2’-O-Me sites (**Fig. 3f, k**). However, van der Waals contacts may not necessarily represent hydrophobic contacts because they also exist with polar residues (**Fig. 3d, j**); also, an O-Me moiety would not be sufficiently polarized to form a hydrogen bond through the methyl group. However, closer inspection shows that without the methylation there would be a gap between polar residues (involving a free 2’-OH group), which would be too far (>4 Å) or inappropriate for a hydrogen bond donor/acceptor system. Hence, creating a van der Waals contact (∼3.5-3.8 Å distance) with a methyl group appears to generate a better stabilization than leaving a gap with no hydrogen bond (e.g. **Fig. 3d, e, h, j, m**).

**Figure 3.**
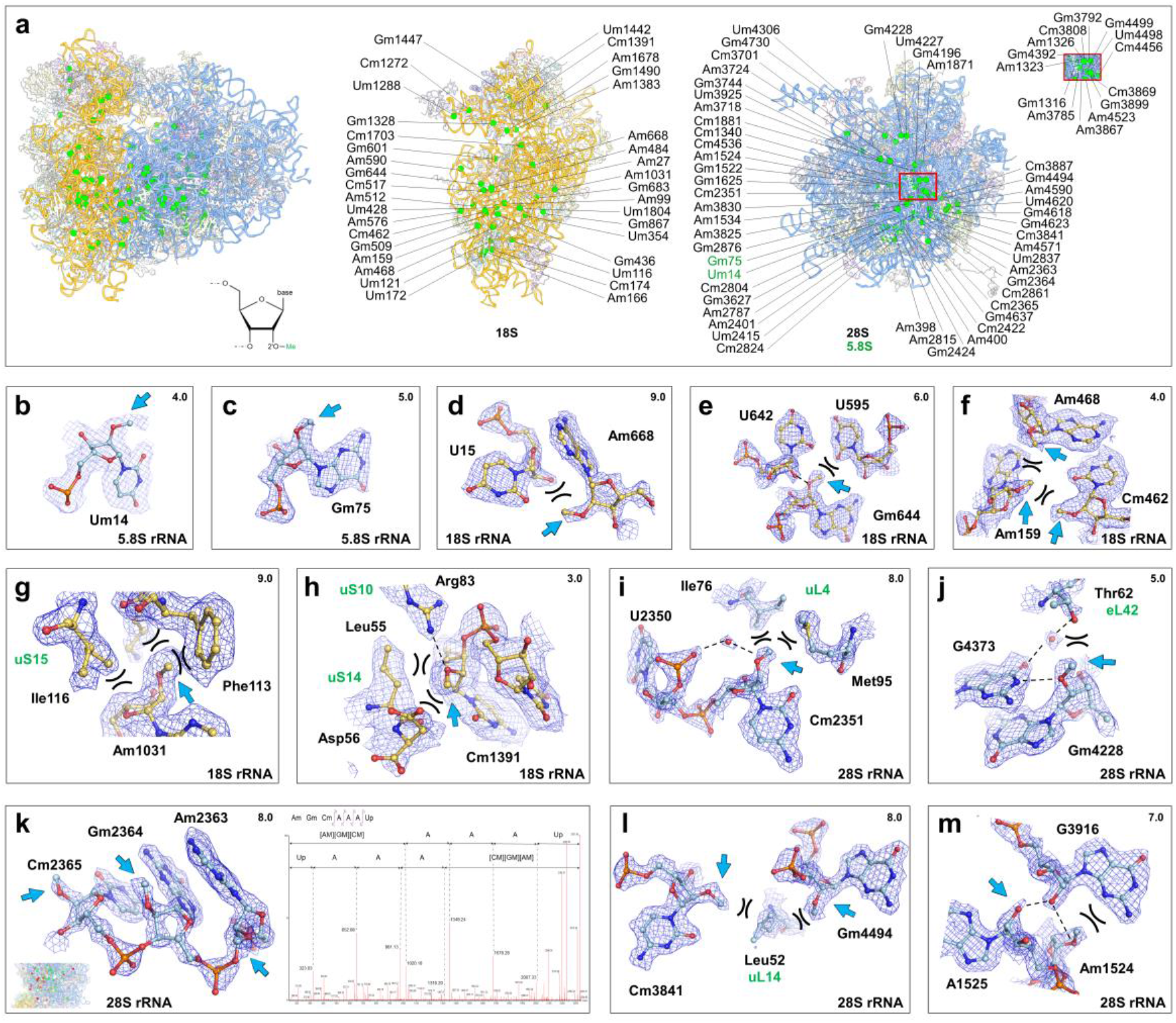
2’-O-methylation sites in the human 80S ribosome. **a** Annotation of 2’-O-Me modified nucleotides on the human 28S, 18S rRNA and 5.8S rRNA of the 40S and 60S ribosomal subunits, respectively. Positions of the 2’-O-methylated nucleotides inside the 80S ribosome are shown as green spheres; 18S rRNA is shown in yellow, 28S rRNA is shown in blue, ribosomal proteins are shown in transparent for clarity. Global front view of the 80S ribosome (left), side view of the 40S subunit (middle) and side view of the 60S subunit (right). Chemically modified nucleotides of the 5.8S rRNA are shown in green. For detailed maps of each site see **Extended Data Figs. 4 & 5**. **b-m** Examples of 2’-O-Me sites. Hydrogen bonds (dotted lines) and van der Waals contacts (black arcs) are indicated. **b-c** 2’-O-methylated nucleotides on 5.8S rRNA (2’-O-methylation are pointed by blue arrows). **d-h** 2’-O-methylated nucleotides and their local environment on the 18S rRNA. **i-m** 2’-O-methylated nucleotides and their local environment on the 28S rRNA. Water molecules are shown as red spheres. **k** 28S rRNA site with 3 successive 2’-O-methylated nucleotides. These modifications are confirmed by mass spectrometry (spectrum on the right), which shows a deconvoluted MS/MS spectrum of ^2363^[Am][Gm][Cm]AAA^2369^Up with fragmentation assignments.

Interestingly, hydrogen bonds can in fact form with the oxygen atom of the 2’-O-Me group, for example with a free ribose 2’-OH group (**Fig. 3e, m**), a polar residue (**Fig. 3h**) or a water molecule (**Fig. 3i**). This is because a methoxy moiety keeps its hydrogen bond acceptor capability due to the 2 free electron pairs. Hence, a methylation removes the hydrogen bond donor activity of a 2’-hydroxy group but still maintains hydrogen bond acceptor activity. Combination of these capabilities thanks to the peculiar chemical properties of a methoxy group can create rather complex contact and interaction patterns, as exemplified by Gm644 and Cm1391 (18S rRNA, **Fig. 3e & h**) or Gm4228 (28S rRNA, **Fig. 3j**). Such series of contacts and interactions involving 2-O-Me moieties contribute to stabilize the conformation of numerous residues, both through van der Waals contacts and - unexpectedly - also hydrogen bonds through the oxygen atom of the methoxy moiety.

### Pseudo-uridine modifications in the human ribosome

As opposed to 2’-O-Me sites which can be readily recognized in a structure, Ψ’s cannot be structurally distinguished from a U due to their isomeric nature in which the C5 and N1 atoms are swapped (**Fig. 4a**). Yet, Ψ sites identified by MS could be confirmed thanks to their specific property of having an exposed hydrophilic N1 position, which translates into the presence of a water molecule (that can be resolved in the high-resolution cryo-EM map), a phosphate group or other polar residue in hydrogen-bonding distance; by comparison, a non-modified U cannot form such interactions in this position. Thanks to this annotation, each site and its structural environment can now be analysed (**Fig. 4**), revealing a characteristic interaction pattern. Ψ’s typically interact via a water molecule with the phosphate group of the same residue (**Fig. 4d, f, h, l**), or with the phosphate group of the n-1 residue (**Fig. 4e, j**) or with both phosphate groups (e.g. 5.8S rRNA, **Fig. 4b, c**), or only with the phosphate group of the n-1 residue (**Fig. 4e, j**). Interactions with the n-2 phosphate (**Fig. 4i**) or with other moieties such as ribose (**Fig. 4g**) or nucleotide bases (**Fig. 4k**, mediated by a water molecule) also exist but are rarer.

**Figure 4.**
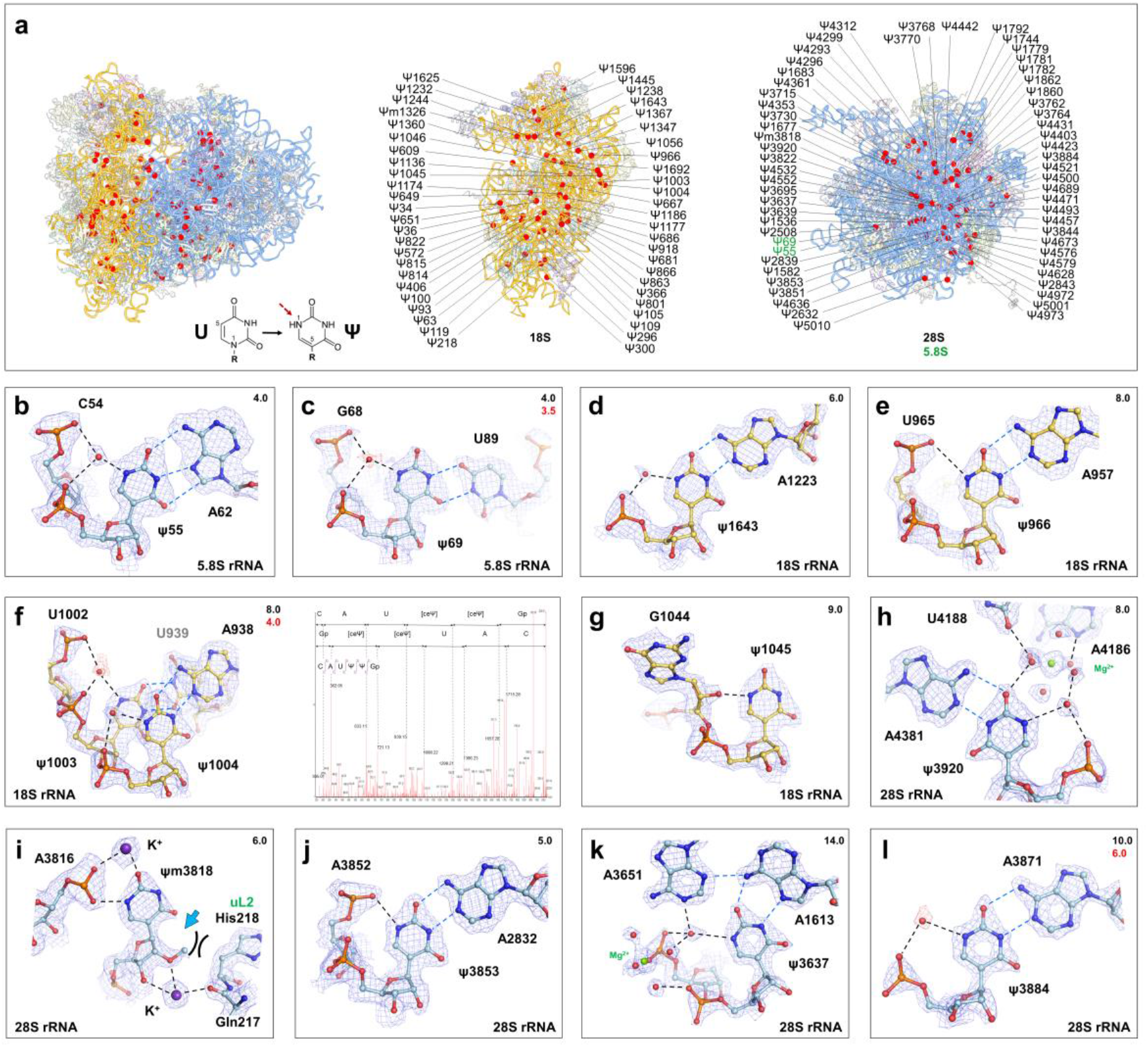
Pseudo-uridines in the human 80S ribosome. **a** Annotation of Ψ-modified nucleotides on the human 28S, 18S rRNA and 5.8S rRNA of the 40S and 60S ribosomal subunits, respectively. Positions of the pseudo-uridine nucleotides inside the 80S ribosome are indicated as red spheres. Chemically modified nucleotides of the 5.8S rRNA are shown in green. For detailed maps and hydrogen bonding distances of each site see **Extended Data Figs. 4 & 5**. **b-l** Examples of a Ψ sites with the characteristic hydrogen-bonding in the N1 position. Water molecules interacting with the N1 atom of pseudo-uridines are shown with red spheres. Hydrogen bonds with water molecules are shown with black dotted lines and hydrogen bonds between bases are shown with blue dotted lines; van der Waals contacts are indicated as black arcs; full visualization of the water molecules at a lower map contour level is indicated in red in panels **c, f, l**. **b-c** Pseudo-uridines on 5.8S rRNA. **d-g** Pseudo-uridines and their local environment on the 18S rRNA. **f** Pseudo-uridines 1003 and 1004 of the 18S rRNA. These were confirmed by mass spectrometry (spectrum on the right), as shown in the deconvoluted MS/MS spectrum of ^1000^CAU[Ψ][Ψ]^1005^Gp with fragmentation assignments; the ceΨ refers to the cyanoethylated adduct. **h-l** Pseudo-uridines and their local environment on the 28S rRNA. Mg^2+^ ions are shown as green spheres and the K^+^ ion interacting with the 2’-O-methylated pseudo-uridine 3818 (Ψm3818; panel **i**) is shown as a purple sphere.

The majority of Ψs (92 of the 109 visible in the structure) have a water molecule bound in the N1 position of the base (**Fig. 4** & **Extended Data Figs. 4 & 5**). These water molecules stabilize the Ψ base conformation because of their interactions with neighbouring polar residues. Many Ψs are in a single-stranded region of the rRNA and/or exhibit hydrogen bonds with a neighbouring nucleotide base. In this structural context, the water molecule-mediated stabilization of the base conformation helps to pre-configure its orientation in a favourable manner for interaction with a neighbouring base, in particular for A-U base pairs (**Fig. 4b, d-f, h, j, k, l**) and non-canonical base pairs such as U-U (**Fig. 4c, f**). The specific stabilization of A-U base pairing through a U-Ψ conversion is particularly frequent and occurs mostly in rRNA helical segments (**Fig. 4**). A-U pairs may benefit from additional hydrogen bonding opportunities because they involve only 2 hydrogen bonds as compared to G-C pairs that have 3. In conclusion, the introduction of a Ψ chemical modification indirectly stabilizes nucleotide base pairing. Consistently, the vast majority of Ψ N1 interactions occurs with rRNA nucleotides but not with amino acids. Overall, Ψ modifications cluster in the 40S ribosomal subunit and the 60S region proximal to the 40S, including at PTC & DC functional centres (**Fig. 4**), for example, the tip of H69 (28S rRNA) carries two universally conserved pseudo-uridines, m^3^Ψ3762 and Ψ3764, located in the tRNA peptidyl site. In conclusion, the presence of the numerous Ψ’s in the structure creates many options for additional hydrogen bonding at the N1 of the modified base, which would otherwise not be available due to the hydrophobic character of the C5 position in a U. As visible from the human rRNA structure, the Ψ can have a local stabilization effect on base pairs under variable configurations (Watson-Crick and Hoogsteen^73^; **Fig. 4b-f,h,j-l**) including base triplets (**Fig. 4k**) and rRNA loops (**Fig. 4g,i**). The creation of additional hydrogen bonds collectively tends to stabilize the macromolecular assembly, consistent with the observation that Ψ-free ribosomes seem to be more flexible structurally^74^.

### Nucleotide base modifications and interfaces

The human rRNA comprises 12 nucleotide base modifications all of which were detected by MS (**Suppl. Data File 1**) and are visible in the structure (**Fig. 5**). The 28S and 18S rRNA’s have 5 and 7 modifications respectively, but none are present in the 5 and 5.8S rRNA’s; the suggested m^5^C4443 (ref. 67) was not found by MS nor seen in the structure, consistent with other studies^12^. The cryo-EM map resolves individual methyl groups on the bases (**Fig. 5b-e, h, k**) or the conformation of the substituent, e.g. the methyl moiety at the N6 position of m^6^A4220 or m^6^A1832 (**Fig. 5b, k**) and the acetyl groups of cytosines C1842 and C1337 (**Fig. 5g, i**). The methyl groups often create stabilizing van der Waals contacts with neighbouring nucleotides (no amino acids involved; **Fig. 5b, c, e, h**) or increase base stacking as the methyl group is co-planar with the base (m^5^C3782, **Fig. 5d, k**). The hypermodified residue m^1^acp^3^Ψ1248 (ref. 75) comprises base methylation and an aminocarboxypropyl (acp) moiety that is more flexible (**Fig. 5f**). Interestingly, its position appears to bridge the 40S head and body regions. Both m^1^acp^3^Ψ1248 and m^5^C3782 are located in the tRNA peptidyl site and are likely to be in contact with the tRNA. In addition, absence of modification in the 1248 position affects 18S rRNA maturation^76^, a mechanism that could make sure that only a properly modified 18S rRNA assembles into an initiation-competent 40S ribosomal subunit. The di-methylated m^6^_2_A1850 and m^6^ A1851 (18S rRNA, **Fig. 5j**) are essential for ribosome assembly^77^. They are located on the 40S ribosomal subunit close to the rotation pivot point next to helix H69 (28S rRNA) corresponding to conformational changes and subunit ratcheting reminiscent of pre-/post-translocation states. m^3^U4530 is located at the PTC next to the entry of the peptide exit channel. Among the two acetylated residues (needed for 40S biogenesis^78^), ac^4^C1842 forms two hydrogen bonds with Arg2 (ribosomal protein eL41), one through its phosphate group and one through the acetyl moiety (**Fig. 5g**), thereby promoting inter-subunit interactions. The other acetylated residue, ac^4^C1337, is located at the entry site of the mRNA channel and might interact with the incoming mRNA (**Fig. 5i**).

**Figure 5.**
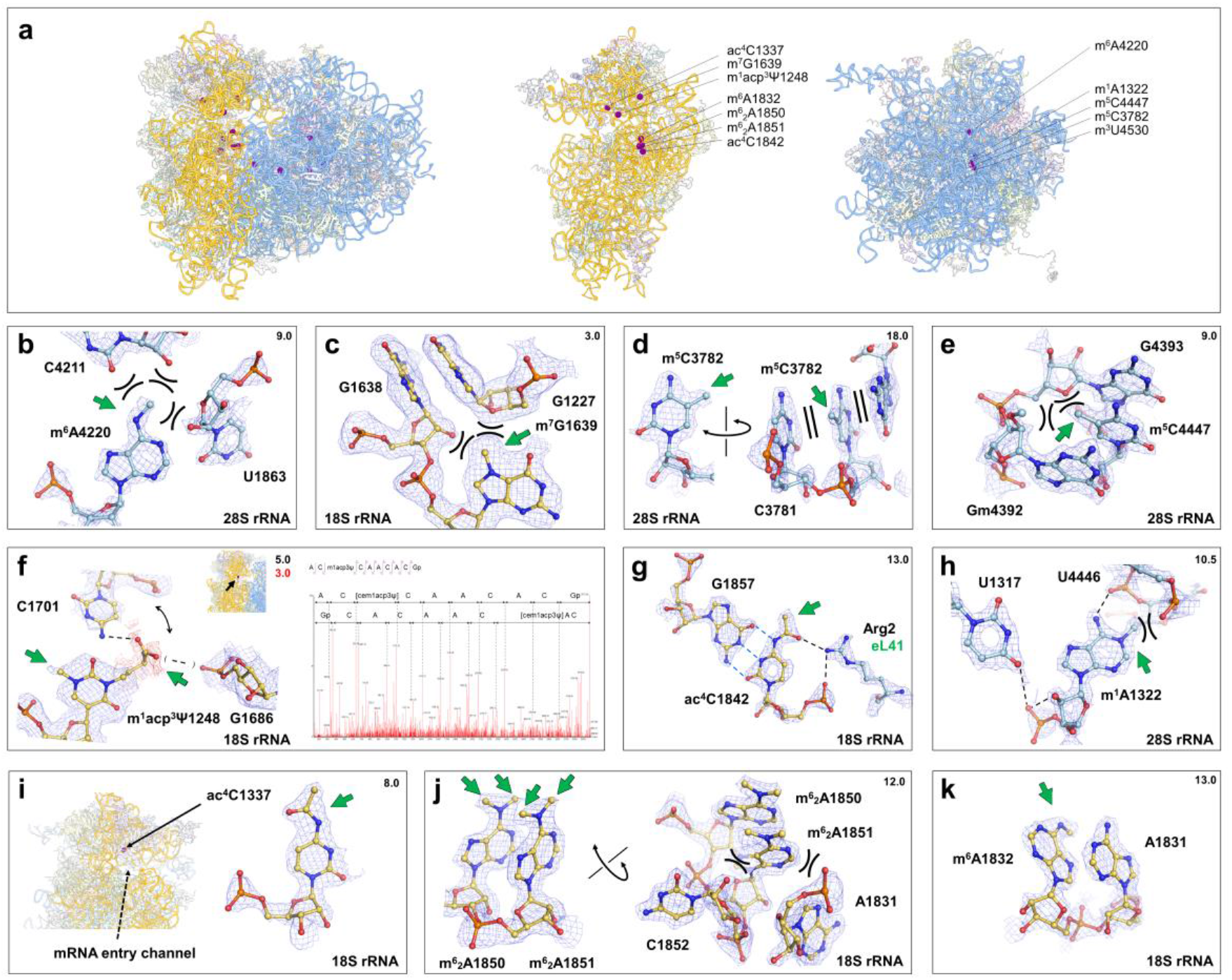
Nucleotide base modifications in the human 80S ribosome. **a** Annotation of nucleotide base modifications (purple spheres) on the human 28S, 18S rRNA and 5.8S rRNA of the 40S and 60S ribosomal subunits, respectively. For detailed maps of each site see **Extended Data Figs. 4 & 5**. **b-k** Various nucleotide base modifications in 18S and 28S rRNA of the human ribosome. Hydrogen bonds with residues are shown with black doted lines and hydrogen bonds between bases are shown with blue doted lines; van der Waals contacts are shown as black arcs. **b** Methylated adenosine 4220 (m^6^A4220) of the 28S rRNA forms numerous van der Waals contacts. **c** Methylated guanosine 4220 (m^7^G1639) of the 18S rRNA forms van der Waals contacts with its environment. **d** Methylated adenosine 4220 (m^5^C3782) of the 28S rRNA forms an increased stacking interaction with its 2 neighboring nucleotide bases (left and right, rotated views). **e** 2’-O-methylated guanosine 4392 (Gm4392) and methylated cytosine C4447 (m^5^C4447) show base stacking that is extended to G4393. **f** Chemically modified nucleotide m1acp3pseudo-uridine and its environment in the P-site. Density of the modification is shown in red at lower contour level to visualize the acp moiety. *Right*, deconvoluted MS/MS spectrum of ^1246^AC[m^1^acp^3^Ψ]CAACAC^1255^Gp from HeLa 18S rRNA with fragmentation assignments. The cem1acp3Ψ refers to the cyanoethylated adduct. **g** Acetylated cytosine 1842 (ac^4^C1842) and its environment. **h** 2’-O-methylated adenosine 1326 (Am1326) and methylated adenosine A1322 (m^1^A1322) with their environment. **i** Acetylated cytosine 1337 (ac^4^C1337) positioned in the 40S ribosomal subunit in proximity of the mRNA channel entry. **j** Pair of dimethylated adenines where stacking is increased and a van der Waals contacts are created with an rRNA loop (left and right, rotated views). **k** m^6^A1832 increases stacking with its neighbor A1831 in the 18S rRNA.

## Discussion

The present high-resolution structure of the human ribosome is a major advance in its observation at the molecular level providing unprecedented insights into its RNA architecture and function. Using latest-generation cryo electron microscope setups and advanced image processing including corrections of higher-order aberrations and focused refinement of separate structural segments has now allowed crossing the 2 Å resolution barrier on the full 80S human ribosome structure. With 1.9, 2.0 and 2.1 Å for the 60S ribosomal subunit and the body and head regions of the 40S ribosomal subunit, respectively, this represents a resolution improvement of 1 Å compared to our previous study^38^ and a subsequent study^79^ [and also slightly better resolved than the isolated 40S subunit (no full ribosome)^80^, which is overall consistent with the present findings]. This type of resolution range is a game changer at the level of observable details (see comparisons in **Extended Data Fig. 3**). This allows resolving unprecedented details on rRNA and ribosomal proteins, chemical modification of the rRNA, water molecules and various ions, but also several post-translational modifications on the ribosomal proteins. It provides the currently best resolved structure of the fully assembled human 80S ribosome including a detailed atomic model of the overall ribosomal RNA and proteins refined to good geometry (even cis-prolines could be built, which is only reasonably possible at such high resolution; **Extended Data Fig. 1**). Thanks to the possibility to visualize water molecules it became possible to annotate pseudo-uridines (Ψ) in the structure; indeed, these uridine isomers cannot be identified in terms of their structure but instead can be indirectly recognized thanks to their resolved N1-associated water molecule.

2’-O-Me, ψ sites and nucleotide base modifications were analysed by advanced mass spectrometry of RNase-induced rRNA fragments. This allowed to confirm the chemical identity of the modification sites and served for cross-validation of the observations made in the structure; for the rare sites located in structurally ill-defined regions at the periphery, the MS data conveniently complete the picture and provide consistency across different biochemical methods used in the past^9,11,12,13,31,69,70,71,72^ (**Suppl. Data File 1**; slight differences could be due to the usage of different cell lines or different chemical identification methods). Together, this combined cryo-EM and MS analysis using the same sample for both methods provides the most complete annotation of chemical modifications in human rRNA. Following our first visualization of chemical modifications in the human ribosome^38^, the present study now resolves the chemical identity of those sites that remained to be characterized and it strongly extends the number of identified chemical modifications in the 3D structure (an additional 91 Ψ’s and 45 2’-O-methylations, confirmed by MS). With the exception of a few sites at the periphery of the structure, all 2’-O-Me, Ψ and base modifications can be seen. In total, 235 modifications were annotated: 220 rRNA modification sites were identified on the entire structure, including 99 2’-O methylations, 109 Ψ and 12 base modifications (the two Ψm’s and m^1^acp^3^Ψ are counted twice due to their hyper-modifications) and an additional 15 sites were seen by MS but not observable in the structure being located at the periphery.

Chemical modifications found by MS and their corresponding sites in the structure are displayed together in **Fig. 6** (see also **Extended Data Figs. 4 & 5** for the detailed cryo-EM map and **Extended Data Fig. 11** for the list of modified fragments detected by MS). Their distribution in the structure is consistent with our observation that an evolutionary extended shell of rRNA modifications exists in eukaryotes^38^. By contrast, the *E. coli* ribosome has much fewer sites (35), of which only 6 are conserved and 3 are chemically different (**Fig. 7**; post-translational modification on the ribosomal protein are also not well conserved, **Fig. 7b**); in *E. coli*, the rRNA modifications are predominantly base modifications (22), contrary to eukaryotes (**Extended Data Figs. 4 & 5** and **Suppl. Data File 1** for the list of modifications in ribosomes from different species). Similar considerations apply also to other bacterial species (e.g. *T. thermophilus*^81^). Even though most rRNA modifications are not conserved *per se*, comparison with the *E. coli* ribosome structure^58^ (**Fig. 7c**) shows that while the residue corresponding to m^6^A1832 (A1500 in *E. coli*) is not methylated its neighboring nucleotide (m^4^Cm1402, corresponding to Cm1703 in human) is methylated instead (**Fig. 7c**). The fact that the methyl moiety is oriented in the same direction in space (possibly for an interacting factor) suggests the existence of a methyl-compensation effect to create a locally hydrophobic environment. The observed base methylation swap between 2 neighbouring residues across human and bacteria (**Fig. 7c**) suggests that the concept of co-evolution of key residues also exists for chemical modifications. Among eukaryotes, rabbit and plant ribosomes^59,82^ comprise chemical modifications similar to human but with some differences (summarized in **Suppl. Data File 1**). The overall distribution of modifications in the human ribosome covers functional centres such as PTC and DC but spreads out further around the tRNA binding sites (**Fig. 6**). Compared to its size (∼1900 nucleotides), the 40S ribosomal subunit comprises a ∼twice higher average occurrence of modification sites (103) than the 60S subunit (132) that has > 5000 nucleotides as compared to the 40S subunit (1870). The strong modification density in the 40S ribosomal subunit appears consistent with a tighter regulation mechanism, in particular during translation initiation and biogenesis. Interestingly, those in the 60S subunit are mainly located in the half oriented toward the 40S ribosomal subunit (**Fig. 6**), hinting at a 40S-associated regulatory role in the 60S subunit. This appears connected with the fact that tRNA binding and translocation and associated conformational changes occur precisely in these core regions of the ribosome, consistent with the observation that modifications may play a regulatory role at molecular level in the translation process^16–27,83^.

**Figure 6.**
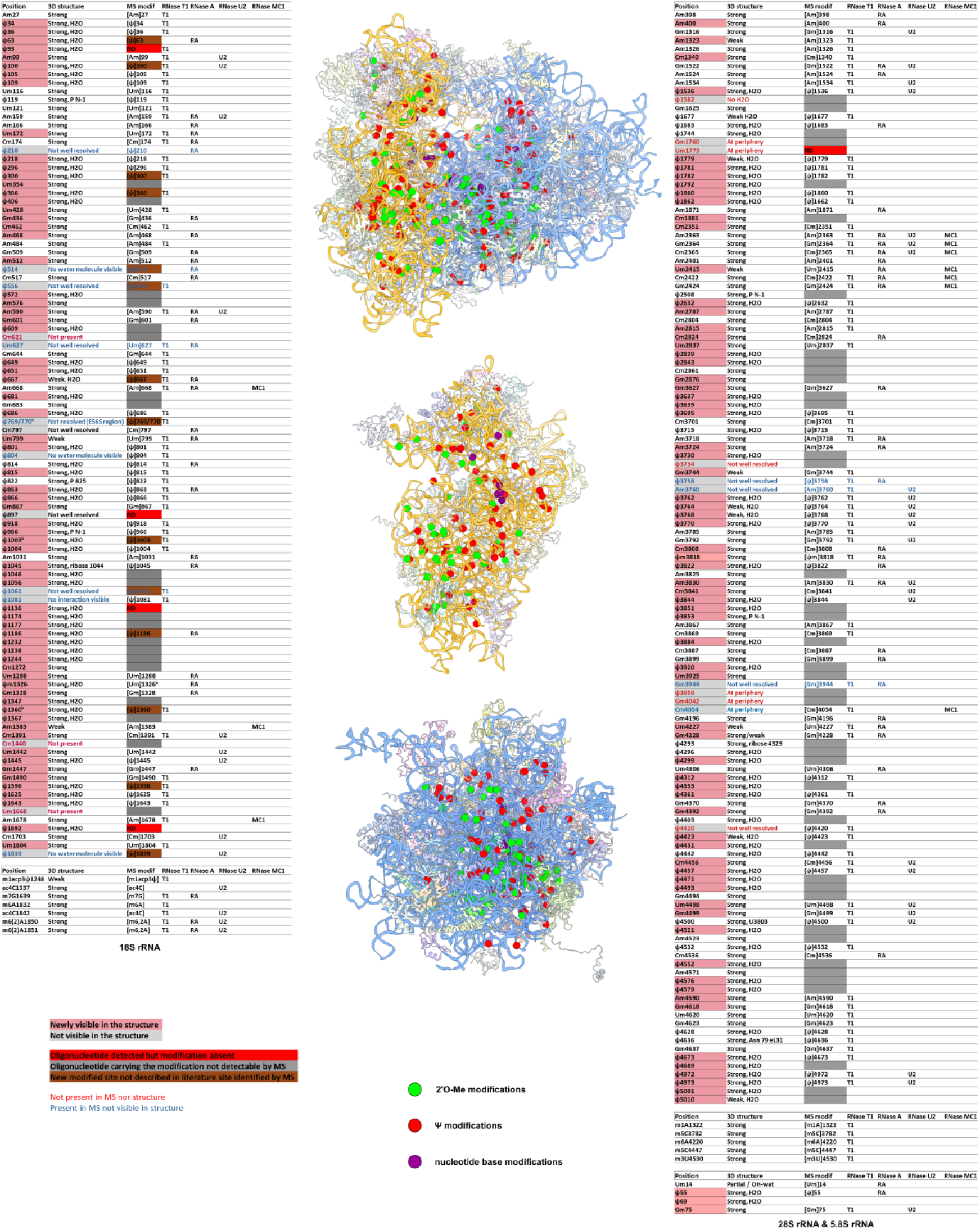
**Complete annotation of chemical modification in the human 80S ribosome through MS and structural analysis** Annotation of 2’-O methylations, Ψ and base modifications by MS (left and right tables for 18S and 28S/5.8S rRNA’s, respectively, and on the 80S human ribosome structure (middle panels). Colours codes for annotations are indicated.

**Figure 7.**
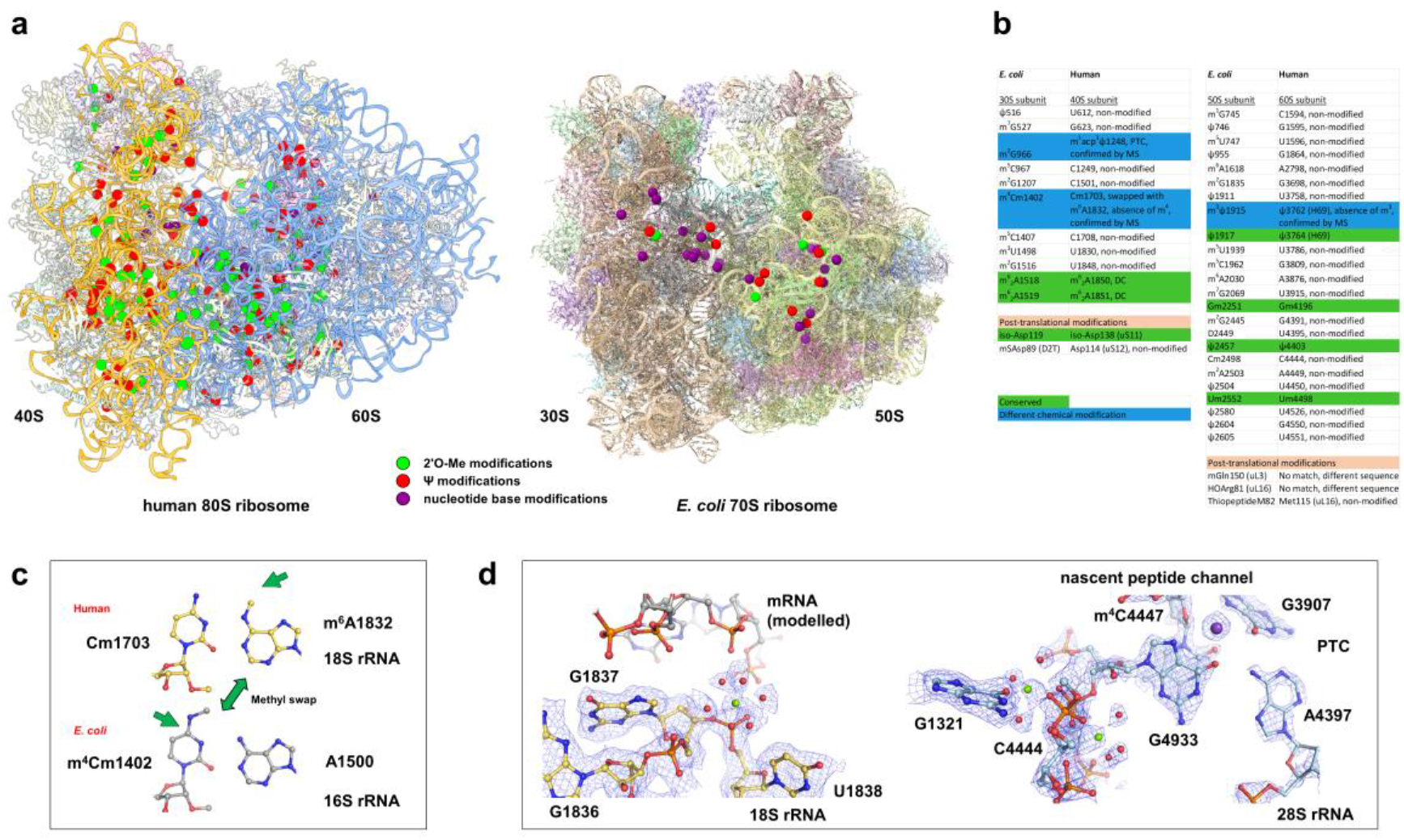
Comparison of chemical modifications and conserved ion sites in the human 80S and *E. coli* 70S ribosome. **a** Annotation of the 235 and 35 2’-O methylations, Ψ and base modifications in human 80S and *E. coli* 70S ribosomes, respectively. Colours codes for annotations are indicated. **b** rRNA chemical modifications in *E. coli* compared to the human ribosome. Only few are conserved (green) and some are chemically different (blue). Post-translational modifications of ribosomal proteins are indicated in orange. **c** Methylation swap between human and *E. coli* ribosomes; while in humans m^6^A1832 is modified, in *E. coli* the neighboring residue is methylated instead, suggesting a co-evolution effect of chemical modifications in this rRNA region. **d** Examples of conserved ion positions at functional sites human versus *E. coli* ribosome. Left, the -1/-2 mRNA region (modelled) on the 40S subunit; right, two hexa-hydrated Mg^2+^ ions (green) and a K^+^ ion (next to G3907, as in Fig. 2) at the PTC and entry to the peptide exit channel.

Due to their opposite hydrophilic versus hydrophobic character, 2’-O-Me and Ψ modifications change the possible hydrogen bonding possibilities and van der Waals contacts. The N1-associated water molecule is positioned within the plane of the nucleotide base and enables new hydrogen bond interactions with neighbouring residues. In particular, one can observe a recurrent stabilization of A-U(Ψ) base pairs through the stabilization of the Ψ base conformation through hydrogen bonding to its own phosphate group and/or to the neighbouring n-1 phosphate group. These specific interaction patterns would not be possible without the introduction of a Ψ modification in these sites. Similarly, the 2’-O-methylation converts a hydrophilic moiety into a partially hydrophobic group, which favours hydrophobic contacts to neighbouring hydrophobic groups, but yet keeping the hydrogen bond acceptor activity on the 2’-O position. The H-bonding peculiarities of 2’-O-Me and Ψ modifications therefore create a versality as compared to non-modified nucleotides, which allows extending the repertoire of interactions directly relevant for molecular recognition mechanisms in the ribosome and in RNA in general, but also beyond in other RNA-nucleoprotein complexes. Finally, the frequent occurrence of Ψ’s allowed discovering that the stabilisation of uridine base conformations through isomerisation and H-bonding helps alleviating the intrinsically less stable A-U base pairing, thus compensating for the reduced amount of H-bonds in an A-U pair (2 H-bonds) as compared to the more stable G-C pair (3 H-bonds). This A-Ψ base pair stabilisation principle sets an interesting concept applicable to RNA in general.

The structure comprises numerous annotated ions and water molecules in their native environment, coordinated by amino acids and nucleotides, hence also clarifying fundamental aspects of chemistry underlying ribosome function. Ions such as Zn^2+^, K^+^ and Mg^2+^ and even their associated water molecules can be localized, revealing the important role that water molecules and ions play at the level of molecular recognition and structural stabilization of the RNA architecture and its interactions with ribosomal proteins and substrates. The characteristic octahedral hydration shell of Mg^2+^ ion comprising 6 water molecules is often seen, but depending on local interactions one or several water molecules can be replaced by coordinating phosphate or ribose moieties of the rRNA or by coordinating amino acids, *i.e.* polar residues are seen to replace precise individual water molecules in Mg^2+^ octahedrons. However, it also turns out that many interactions are mediated by water molecules; for example, phosphate groups do not necessarily interact directly with Mg^2+^ ions in contrast to what could be commonly thought. The Mg^2+^ ion can adopt various configurations with varying numbers of associated water molecules. Mg^2+^ ions and their associated water molecules contribute to bridging structural elements within RNA and at protein-RNA interfaces and thereby enable ion-and water molecule mediated interactions of structural elements that would otherwise be too far from each other to interact. Zn^2+^ ions which are found mostly in the eukaryote-specific ribosomal proteins provide extra stabilization as these are surface-exposed proteins on the ribosome; this striking feature hints at an evolutionary predilection of recruiting Zn^2+^-binding proteins to the surface of the translation machinery.

Finally, hydrogen bonds within Mg^2+^ octahedrons or K^+^ ions often extend to nearby water molecules to form a network. This hydration shell is not only visible at the surface of the ribosome (as classically found for many protein structures in the PDB) but also in many cavities inside. Locally and collectively, the numerous ion and water molecule interactions with rRNA and ribosomal proteins contribute to stabilize the 3D structure by holding structural elements of the ribosome together, yet providing a sort of flexible, dynamic network that can fluctuate and adapt along with conformational changes of the ribosome during the translation process. Interestingly, some Mg^2+^ octahedrons and K^+^ ions are found in the same position in human, yeast, plant and *E. coli* ribosomes suggesting a conserved role, for example in the PTC (**Fig. 7d**). The existence of conserved sites for Mg^2+^ and K^+^ ions throughout the ribosome indicates a certain requirement of rRNA for ion binding in specific sites to fold into its active state. In a general way, the manner by which ions such as hexa-hydrated Mg^2+^ clusters and K^+^ interact within protein-RNA assemblies is of broader interest for the analysis of macromolecular complexes, but also from the chemistry point of view regarding coordination chemistry and catalysis.

Taken together, the integrated structural analysis cross-validated with extensive MS-analysis has fostered a unique synergy between structural and biochemical analysis of rRNA chemical modifications allowing a comprehensive analysis of their three-dimensional environment at the molecular level. This study represents currently the largest characterisation of a single set of RNA modifications in one single assembly (these structure & MS cross-validated data have now been integrated into the latest version of the Modomics database^84^). Thanks to our rRNA sequencing we could now identify which rRNA variant is present in HeLa cells, which is important considering that HeLa cells are widely used and that differential variants exist in cancer cells. Being on the human system the findings are relevant for human health, but also for understanding translation mechanism in eukaryotes in general. As a mature 80S ribosome assembly the structure gives unique insights into each ribosomal subunit and their joint interface at which numerous ions and chemical modifications are located. This will help guiding future functional analysis of modification enzymes and help understanding human ribosome function in the healthy context and diseases because modifications (which are introduced during biogenesis) are involved in translation errors, accuracy loss and antibiotic activity changes. The present work thus paves the way towards functional assays regarding specific modification sites of interest to address the molecular mechanism underlying dysregulation and human diseases considering the growing implications of the human ribosome as a cancer target^32,33,36,37^. The structure will serve as a reference for the structure-function analysis of various functional complexes with translation factors, mRNA and tRNA and as a high-resolution template for molecular dynamics simulation calculations, modelling and drug design.

## Supporting information

List of sites analysed by MS and cryo-EM and the corresponding annotations

## Acknowledgments

We thank the TFS and Jeol companies for kindly making their microscope setups available for test data collection, which was performed by Evgeniya Pechnikova (TFS, Eindhoven, NL) and Fumiaki Makino (on behalf of Jeol, Osaka, Japan). We acknowledge Jonathan Michalon, Mathieu Schaeffer and Sasha Ballet for IT support and the IGBMC cell culture facilities for HeLa cell production, members of the integrated structural biology platform at CBI for support and the late Jean-François Ménétret for his constant interest. A.L. and P.W. thank Béatrice Chane-Woon-Ming for the implementation of the MassSpec-Toolkit for RNAs software. This work was supported by CNRS, Association pour la Recherche sur le Cancer (ARC), Institut National du Cancer (INCa_16099), the Fondation pour la Recherche Médicale (FRM & FDT202304016898), Ligue nationale contre le cancer (Ligue), Agence Nationale pour la Recherche (ANR) and USIAS of the University of Strasbourg (USIAS-2018-012). This work of the Interdisciplinary Thematic Institute IMCBio, as part of the ITI 2021-2028 program of the University of Strasbourg, CNRS and Inserm, was supported by IdEx Unistra (ANR-10-IDEX-0002) and by SFRI-STRAT’US project (ANR 20-SFRI-0012), EUR IMCBio (ANR-17-EURE-0023) under the framework of the France 2030 program and LabexNetRNA (ANR-10-LABX-0036_NETRNA) administered by ANR and by the epiRNA funding from the Region Grand Est. The electron microscope facility was supported by the Region Grand Est, FEDER, the French Infrastructure for Integrated Structural Biology (FRISBI) ANR-10-INSB-05-01 / France 2030 program, Instruct-ERIC and iNEXT-Discovery.

Correspondence and requests for materials should be addressed to B.P.K. Manuscript distributed under a Creative Commons Attribution 4.0 International CC-BY licence https://creativecommons.org/licenses/by/4.0/. The authors declare no competing financial interests.

## Author Contributions

I.H., L.F. & S.H. performed sample preparation, L.F., S.H. & O.vL. image processing, A.L. & P. W. MS analysis and annotations, S.N.T.D. & O.vL. rRNA sequencing, C.B., L.F., S.H., O.vL. & B.P.K. structure refinement & model building, structural analysis and annotations. All authors analysed the data. B.P.K supervised the project and wrote the manuscript with input from all authors.

## Author Information

The authors declare no competing financial interests. Correspondence and requests for materials should be addressed to B.P.K.

## Methods

### Complex formation and cryo-EM data processing

Human 80S ribosomes were prepared from HeLa cells (certified as free of Mycoplasma using a polymerase chain reaction test) as described earlier^51^. Cryo-EM grids (carbon-coated Quantifoil R 2/2 cupper grids, 300 mesh) were plunge-frozen using a Vitrobot IV (Thermo Fisher Scientific) operating at 95% humidity and 10°C.

Three data sets on the human 80S ribosome were used: data set 1 was collected on a Titan Krios G4 (equipped with a Selectris X energy filter and Falcon 4 camera; TFS factory in Eindhoven) with a pixel size of 0.72 Å as EER movie (total dose 40 electron / Å^2^) and 2 & 3 were collected on a CRYO ARM 300 (equipped with an Omega energy filter, Jeol, and a K3 camera, Gatan; RIKEN Osaka cryo-EM facility) at a pixel size of 0.82 Å as compressed .tif movies, respectively, both microscopes operating at 300 kV^85^. The three data sets were first processed separately (see also **Extended Data Figs. 1 & 2**), including movie alignment using Relion’s motion correction algorithm^86^ based on MotionCor2^87^, CTF estimation using GCTF software^88^ implemented in Relion, particle picking using crYOLO^89^. All further steps were done in Relion^86^. From the initially obtained particles sets (634 158 particles were obtained for data set 1, 484 393 for data set 2 and 149 674 particles for data set 3) 199 371, 229 233 and 105 596 particles were kept after 2D & 3D classifications. The full-sized particles were refined in 3D followed by CTF correcting for anisotropic magnification, beam tilt, trefoil, 4^th^ order aberrations, defocus and astigmatism followed by Bayesian polishing. Particles from the three data sets were joined as distinct optics groups (merged data set) and refined together in 3D followed by a 3D global classification to 5 classes without alignment (**Extended Data Fig. 2**). The largest class was kept (382 016 particles) and refined to 2.22 Å resolution. From this map three masks containing 60S large ribosomal subunit, 40S small ribosomal subunit body 40S small ribosomal subunit head and E-site tRNA were generated and extended by 12 pixels with a soft edge of 7 pixels with the *relion_mask_create* tool. These masks were used for focused refinements and the multi body refinement^41–50^. A final resolution of 1.93 Å was obtained for the 60S large ribosomal subunit, 2.06 Å for the 40S small ribosomal subunit body and 2.14 Å for the 40S small ribosomal subunit head as estimated from Fourier shell correlation calculations^90–93^ with the 0.143 cut-off criterion. Using partial signal subtraction on each body further improved the map (**Extended Data Fig. 3**), resolving for example a whole set of additional hexa-hydrated Mg^2+^ ions (particularly in the 40S ribosomal subunit). The post-processing procedure implemented in Relion^86^ was applied to the final maps for masking, B-factor sharpening (-36.46, - 37.99, -49.39 Å^2^ for the 60S, 40S body and 40S head regions, respectively) using masks prepared from 15 Å lowpass filtered maps, that were shrunk by 3 pixels using a soft edge of 9 pixels. Local resolution estimation was done using the local resolution job of Relion.

### Atomic model building & refinement and localisation of rRNA modifications

The maps of the three sub-regions were combined into a single 80S map and an atomic model of the human ribosome (PDB: 6qzp^38^) was placed as a rigid body into the map using Chimera^93^. All positions for chemical modifications of the rRNA detected by the present MS analysis as well as previously reported rRNA modifications^38,95,96^ were manually cross-checked for presence in the structure, as summarized in **Suppl. Data File 1**. Chemical modifications were identified based on the presence of significant additional densities in the 2’-OH position (indicated by cyan arrows in the figures) and nucleotide bases. The map contour level was adjusted individually as indicated by the local root mean square deviation from the mean value (similar to sigma levels) in each figure corner. Mg^2+^, K^+^ and Zn^2+^ ions were annotated in the structure based on their characteristic coordination patterns and coordination distances^97^. Typical distances found for water molecules and ions are 2.3 Å for Zn^2+^, 2.1 Å within the octahedron for Mg^2+^ and 2.6-3.2 Å range for K^+^, respectively, consistent with values known from chemistry and in proteins^63^. Water-coordinated Mg^2+^ and modified amino acids and nucleotide templates were drawn using JLigand^98^ and their respective restraints for the atomic model fitting and refinements were generated. The model was subsequently refined in Phenix^99^ using restrained parameter refinement protocols (real space refinement, positional refinement, grouped B-factor refinement and simulated annealing as described^36,52^. Finally, water molecules were picked using an automated procedure that combines density search and coordination with hydrophilic residues considering potential hydrogen bonds using SWIM^63^ and Phenix^56,64^ software; we used the peak search implemented in Chimera using the segmentation-guided water and ion modelling (SWIM) through the Segger software^63^ plugin to the UCSF Chimera^100^ software. Water molecule annotation was done after having annotated (i) Zn^2+^ ions in their tetrahedral environment coordinated by cysteine residues, (ii) Mg^2+^ ions with their associated water molecules in an octahedral coordination, (iii) K^+^ ions and (iv) water molecules at the N1 position of Ψ’s (if present, sometimes visible at lower contour level of the cryo-EM map). For regions with weaker cryo-EM density it cannot be excluded that some of the annotated water molecules are ions, but we refrained from annotating these else than as water molecules. The overall structure comprises 5783 nucleotides and 11440 amino acids from the 4 rRNAs and 80 ribosomal proteins, respectively (see statistics in **Extended Data Fig. 1**). Images were prepared with ChimeraX 1.0^100^ and Pymol^101^.

### Mass spectrometry analysis

#### Chemicals and Reagents

Chemicals used were of analytical grade or high purity unless indicated otherwise. Water used to prepare sample solutions or buffers was obtained from Merck Millipore. Ammonium acetate (NH_4_OAc), triethylamine (TEA), ZipTip®C18, Tri-Reagent® and acrylonitrile were purchased from Sigma-Aldrich. 1,1,1,3,3,3-hexafluoropropan-2-ol (HFIP) was purchased from Honeywell. RNase T_1_, RNase A and RNase H were purchased from ThermoFisher Scientific. RNase U_2_ as well as RNase MC1 were homemade in 100 mM ammonium acetate (pH is not adjusted). Oligonucleotides used for RNA-DNA hybridization were synthesized by Eurofins Genomics.

#### Culture of HeLa cells and rRNA isolation

HeLa cells were obtained from a 10 L culture as described previously^36,38,51,52^. Total RNA was extracted from the cells using Tri-Reagent following the manufacturer procedure. Next, the 28S and the 18S rRNA were isolated using SEC (Superose 6 Increase 10/300 GL column) by isocratic elution with 300 mM NH_4_OAc at 0.5 mL/min on an ӒKTA purifier system (General Electric Healthcare). The 5.8S and 5S were isolated from the total RNA extract by 10% polyacrylamide gel. Gels were stained by ethidium bromide and the bands containing the rRNAs were visualized and excised under UV light. Gels bands were dried under vacuum.

#### Cyanoethylation of pseudo-uridines

Purified 28S and 18S rRNA samples were subjected to cyanoethylation following the procedure as described with few differences^55,102^. The reaction was carried out with 10 µL of purified rRNA sample, plus 150 µL of 41% EtOH / 1.1 M TEAA (Triethylammonium acetate) (pH 8.5) and 10 µL of acrylonitrile. Specific labeling of pseudo-uridines was performed for 2 h at 70°C. Cyanoethylated rRNA samples were cooled down and precipitated by adding 40 µL 3M NH_4_OAc, 40 µL H_2_O and 600 µL EtOH 100% and stored at -20°C for at least 2h. Samples were centrifugated at 12 000 g for 10 min at 4°C. The supernatant was removed and the samples were washed with 75% EtOH. Pellets were dissolved in 300 mM NH_4_OAc.

For the 5S and 5.8S rRNAs, the cyanoethylations were achieved in gel. In a tube containing the dried gel, 30 µL of 41% EtOH / 1.1 M TEAA (pH 8.5) plus 4 µL of acrylonitrile were added. Labelling was carried out at 70°C for 2 h. After derivatization, supernatant was removed and the gel were washed 3 times with 200 mM NH_4_OAc. Pieces of gel were then dried under vacuum.

The modification m^1^acp^3^ψ1248 (18S rRNA) reveals an extra mass corresponding to the addition of a cyanoethyl group. Considering that the only reactive nitrogen atom available is located on the modification, this indicates the formation of a cyanoethylated acp3 adduct due to the chemical treatment, which to our knowledge has not been observed before.

#### RNase H cleavage and gel electrophoresis

The specific cleavage of rRNA was done as described^53,55^. Prior RNase H cleavage, rRNA (≈ 10 µg) and oligos (≈ 10 µg) were mixed and heated at 80°C for 2 min followed by a cooling down to 50°C for DNA-RNA hybridization (see list of oligos in **Extended Data Fig. 10**). To obtain a specific cleavage, a chimeric oligo was also used (see spectrum in **Extended Data Fig. 9**). DNA-RNA duplexes were digested at 50 °C for 30 min by adding 1 µL of 10X RNase H buffer then 2.5 µL of 0.5 U/µL RNase H. After RNase H cleavage, the fragments of interest were isolated by 10% denaturing polyacrylamide gel. Gels were run at 16W for about 4 h. After migration, gels were stained by ethidium bromide and the bands containing the fragments of interest were visualized and excised under UV light. Bands were dried under vacuum. RNA & DNA oligos used for mass spectrometry analysis are listed in **Extended Data Fig. 10**. 18S, 28S, 5.8S & 5S rRNA fragments with 2’-O-Me’s, ψ’s & nucleotide base modifications that were analyzed by mass spectrometry analysis are summarized in **Extended Data Fig. 11**.

#### RNA digestion and NanoLC-MS/MS of RNA oligonucleotides

Dried pieces of gel containing RNA were digested by adding 20 µL of RNase T_1_ (1/1000) or 20 µL of RNase A (1/1000). Samples were then incubated at 55°C for 2h. In-gel digestions by RNase U_2_ were carried out following previously described procedures^103,104^. Briefly, 20 µL of 0.1 µg/µL U of RNase U_2_ in 200 mM NH_4_OAc (pH 5.3) samples were incubated at 55°C for 30 min. The MC1 digestions were carried out in solution for the 28S and 18S rRNAs because gel digestion did not work. Briefly, 20 µL of 0.1 µg/µL U of RNase MC1 in 100 mM NH_4_OAc was added to 10 µg of rRNA. MC1 digestions were achieved in 37°C for 1 h. After digestion, the samples were desalted with ZipTip C18 using 200 mM NH_4_OAc. Eluates were dried under vacuum and pellets containing RNase digestion products were resuspended in 3 µL of milliQ water prior injection. Separation of products were achieved on an Acquity peptide BEH C18 column (130 Å, 1.7 µm, 75 µm × 200 mm) using a nanoAcquity system (Waters). The analysis was performed with an injection volume of 3 µL. To avoid RNase contamination, the chromatographic system was thoroughly washed when analyzing samples digested with another RNase. The column was equilibrated with buffer A containing 7.5 mM TEAA, 7.0 mM TEA and 200 mM HFIP at a flow rate of 300 nL/min. Oligonucleotides were separated using a gradient from 15% to 35% of buffer B (100% methanol) for 2 min followed by an increase of buffer B to 50% in 20 min. Column oven temperature was set at 65°C. Detection by MS and MS/MS were achieved using SYNAPT G2-S from Waters Corporation. The source setting used were: polarity mode: Negative ion; Capillary voltage: 2.6 kV; Sample cone voltage: 30 V; Source temperature: 130°C. The samples were analyzed over an m/z range from 550 to 1600 for the full scan, followed by fast data direct acquisition scan (Fast DDA). Collision-induced dissociation (CID) experiments were achieved using Argon gaz. Data were analyzed using the MassLynx software^105^. Oligonucleotides sequences were confirmed by MS/MS deconvoluted spectra by following c or/and y ion series.

### Sequencing of rRNA

To clarify sequence inconsistencies in data banks we performed sequencing of the 28S, 18S, 5.8S and 5S of the rRNAs to unify the annotation of nucleotides and chemical modifications. 5S, 5.8S, 18S and 28S rRNA samples purified from HeLa cell extract were reverse transcribed using transcriptor reverse transcriptase (Roche) and amplified by PCR using Pfusion PCR polymerase (NEB) with specific primers or random hexamer primers. Here, total RNA samples were used for 5S and 5.8S rRNA. For the 18S and 28S rRNA the RNA was purified as described in the MS section. The rRNA sequence was annotated with rRNA modifications found in the structure and detected by MS (**Extended Data Fig. 13**). Primers used for PCR are listed in (**Extended Data Fig. 14**). The PCR products were sent for sequencing (Eurofins Genomics). Structure, MS and rRNA sequencing allowed a mutual validation and some sequence corrections as compared to the genomic sequence found in data banks (e.g. position 4910 in the 28S rRNA was found to be adenine instead of guanine, visible also in the structure; **Extended Data Fig. 12**).

### Data availability

Atomic coordinates and cryo-EM maps have been deposited in the Protein Data Bank and EMDB under accession codes 8QOI and EMD-18539 (full 80S ribosome), EMD-18812, EMD-18813 and EMD-18814 for the focused refinements of the 60S ribosomal subunit, the 40S ribosomal subunit body and head regions, respectively, and EMD-18815 for the last global refinement of the 80S ribosome before the focused refinements. Mass spectrometry data have been deposited in the ProteomeXchange Consortium via the PRIDE data base http://www.ebi.ac.uk/pride with the dataset identifiers PXD046739, PXD046743, PXD046744, PXD046747 for the 28S, 18S, 5.8S and 5S rRNA’s, respectively. All other materials such as cells are available upon reasonable request.

**Legend to Supplementary Data File 1**

Complete list of sites analysed by MS and cryo-EM and the corresponding annotations, including previously reported rRNA modifications^38,94,95^. RNAses that were used to identify the respective modifications are indicated. Chemical modifications from *E. coli*, rabbit and plant ribosome structures^57,58,59,81^ are included for comparison.

