## Extended Figures for "Structure of the human 80S ribosome at 1.9 Å resolution – the molecular role of chemical modifications and ions in RNA"

**to**

**Table 1**

| **Image processing** | **Data set 1** | **Data set 2** | **Data set 3** | **Merge data set** |
|---|---|---|---|---|
| Total number of extracted particles | 634158 | 484393 | 149674 | 534190 |
| Final number of particles used | 199371 | 229233 | 105596 | 382016 |
| Pixel size (Å) | 0.72 | 0.82 | 0.82 | 0.82 |
| Box size (pixels) | 640 | 512 | 512 | 512 |
| Symmetry | C1 | C1 | C1 | C1 |
| Map sharpening B-factor (Å$^2$) | -37.3 | -41.1 | -39.4 | -36.5 (60S)<br>-38.0 (40S body)<br>-40.4 (40S head) |
| Global map resolution (Å) | 2.1 | 2.2 | 2.0 | 1.9; 2.0; 2.1 |

**Table 2**

| **Atomic model refinement** | |
|---|---|
| Model composition<br>Chains<br>Atoms<br>Residues (a.a. / nucleotides)<br>Water molecules<br>Ions<br>$Zn^{2+}$<br>$Mg^{2+}$ (octahedral)<br>$K^{+}$ | <br>91<br>233497<br>11440; 5783<br>16411<br>410<br>9<br>294<br>107 |
| Bonds (rmsd)<br>Length (Å)<br>Angles (°) | <br>0.004<br>0.801 |
| *MolProbity* score | 1.87 |
| Clash score | 8.27 |
| Ramachandran plot (%)<br>Favoured<br>Allowed<br>Outliers | <br>96.55<br>3.45<br>0.11 |
| Rotamer outliers (%) | 1.74 |
| Cβ outliers (%) | 0.00 |
| Peptide plane (%)<br>Cis proline / general<br>Twisted proline / general | <br>1.7 / 0.1<br>0.0 / 0.0 |
| ADP B-factors (mean)<br>Protein<br>Nucleotides<br>Ligand<br>Water molecules | <br>23.8<br>32.1<br>15.9<br>16.1 |
| Resolution (Å)<br>Range used during refinement<br>$d_{99}$ (full/half1/half2)<br>$d_{model}$<br>$d_{FSC\text{-}model(0,0.142,0.5)}$ | masked; unmasked<br>1.9; 1.9<br>1.7; 1.7<br>1.9; 1.9<br>1.6 / 1.8 / 2.0; 1.6 / 1.8 / 2.1 |
| Model versus data<br>CC (mask)<br>CC (Box)<br>CC (peaks)<br>CC (volume)<br>Mean CC for ligands | <br>0.80<br>0.78<br>0.75<br>0.79<br>0.60 |

**Data collection and refinement statistics**

**Extended Data Fig. 1**

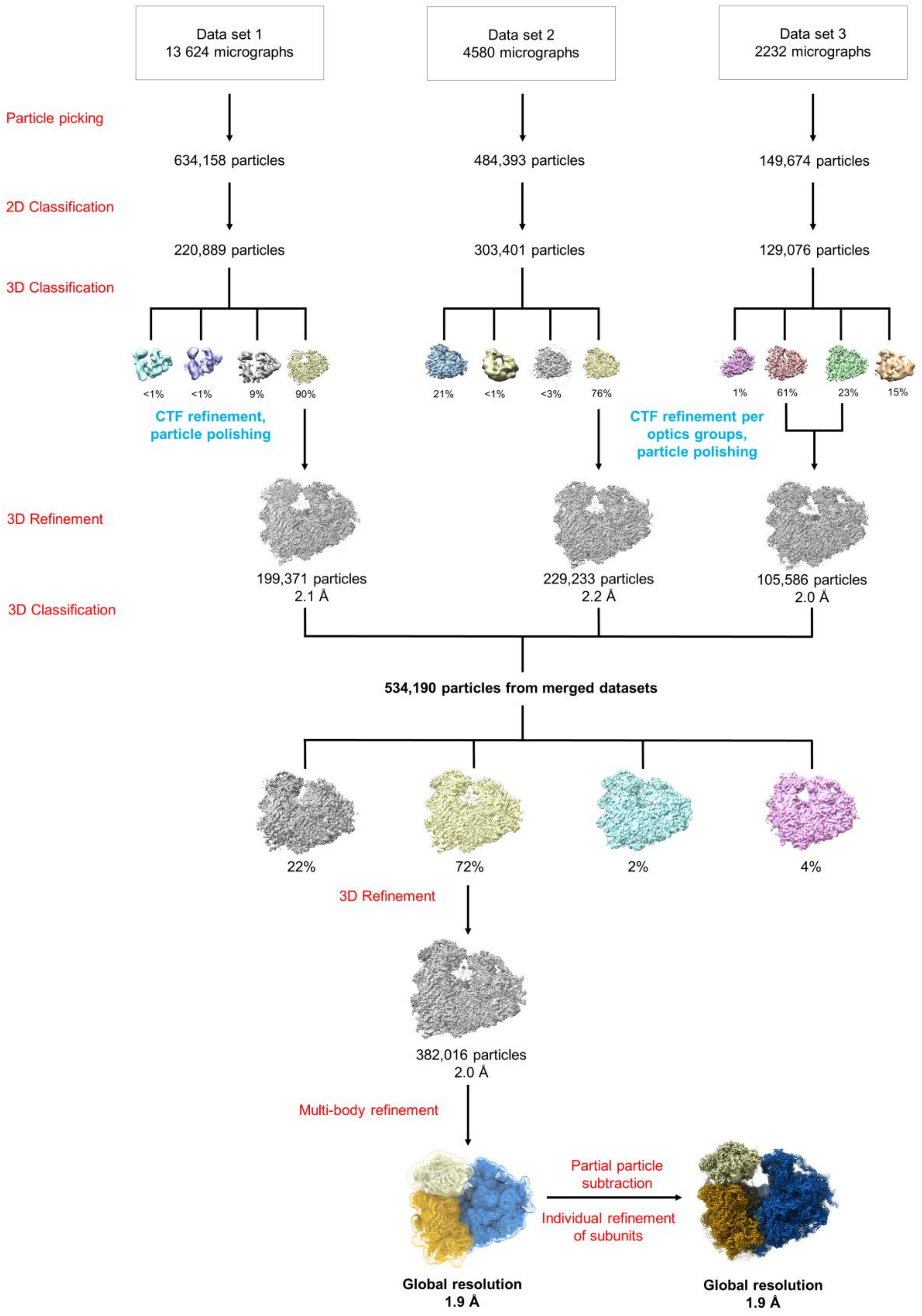

**Particle sorting scheme**

**Extended Data Fig. 2**

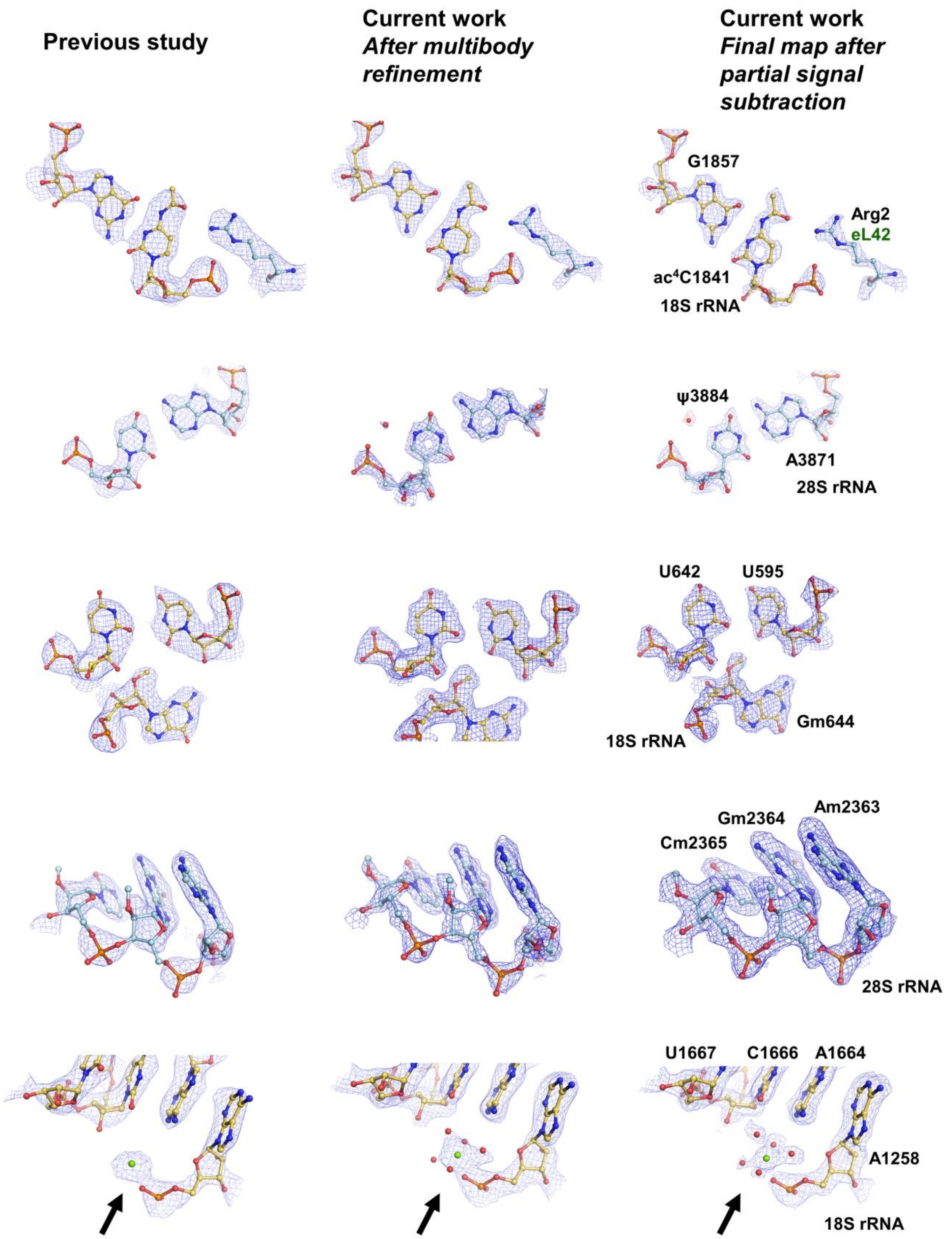

**Resolution improvement of features in the cryo-EM map (present work compared to our previous study[38])**

**Extended Data Fig. 3**

**18S rRNA – 2'-O-methylations**

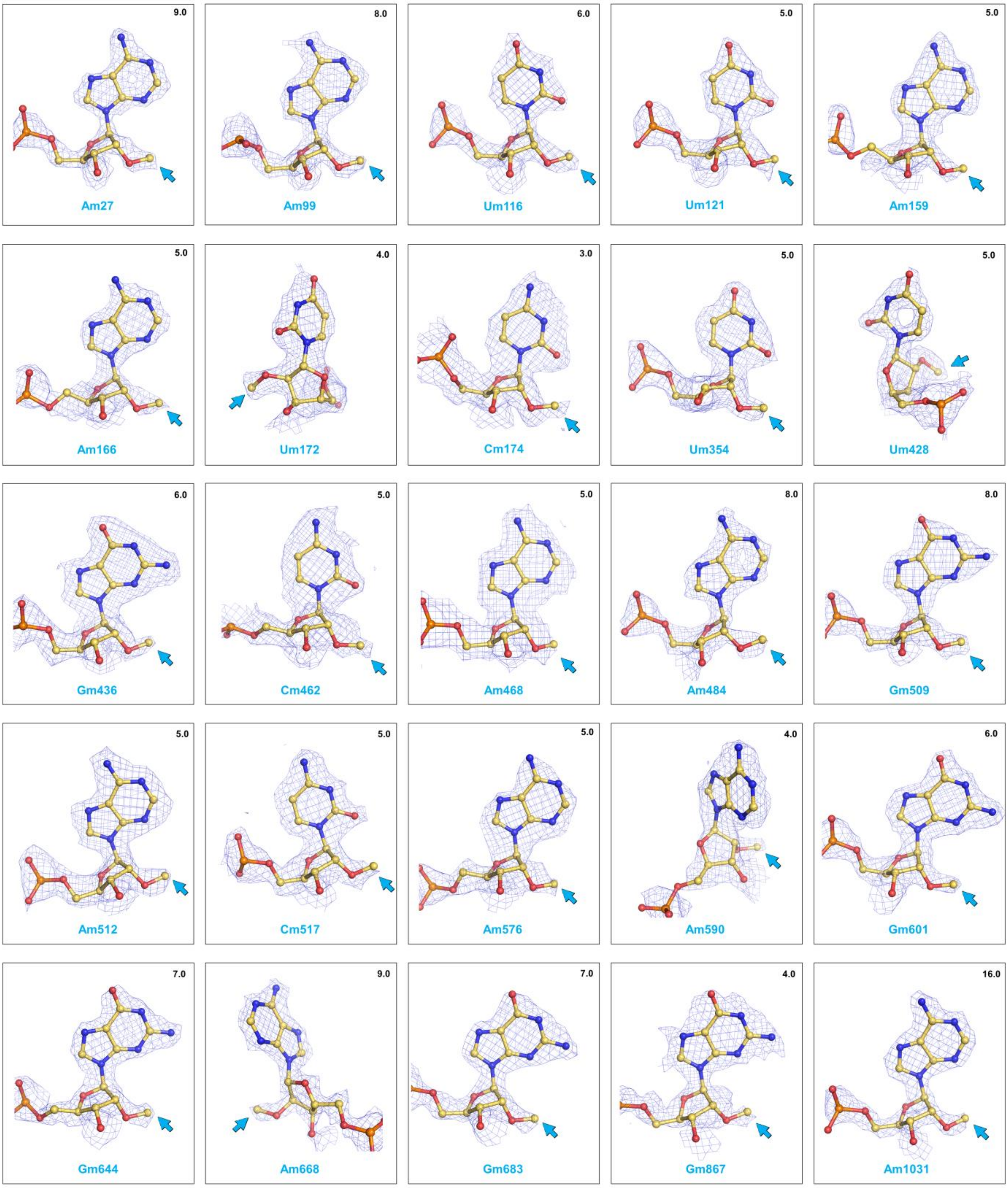

18S rRNA – 2’-O-methylations (cont.)

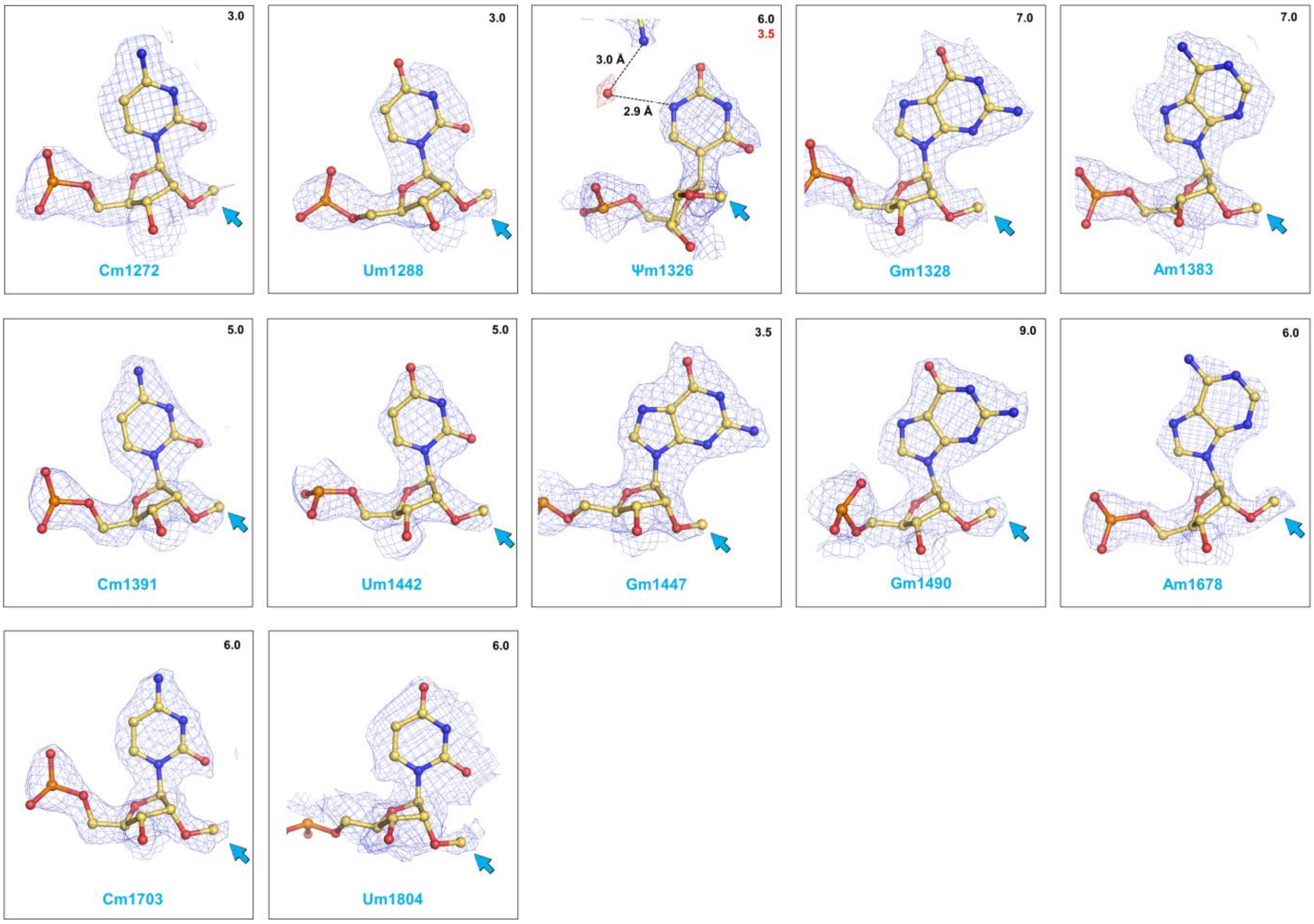

18S rRNA – pseudo-uridines

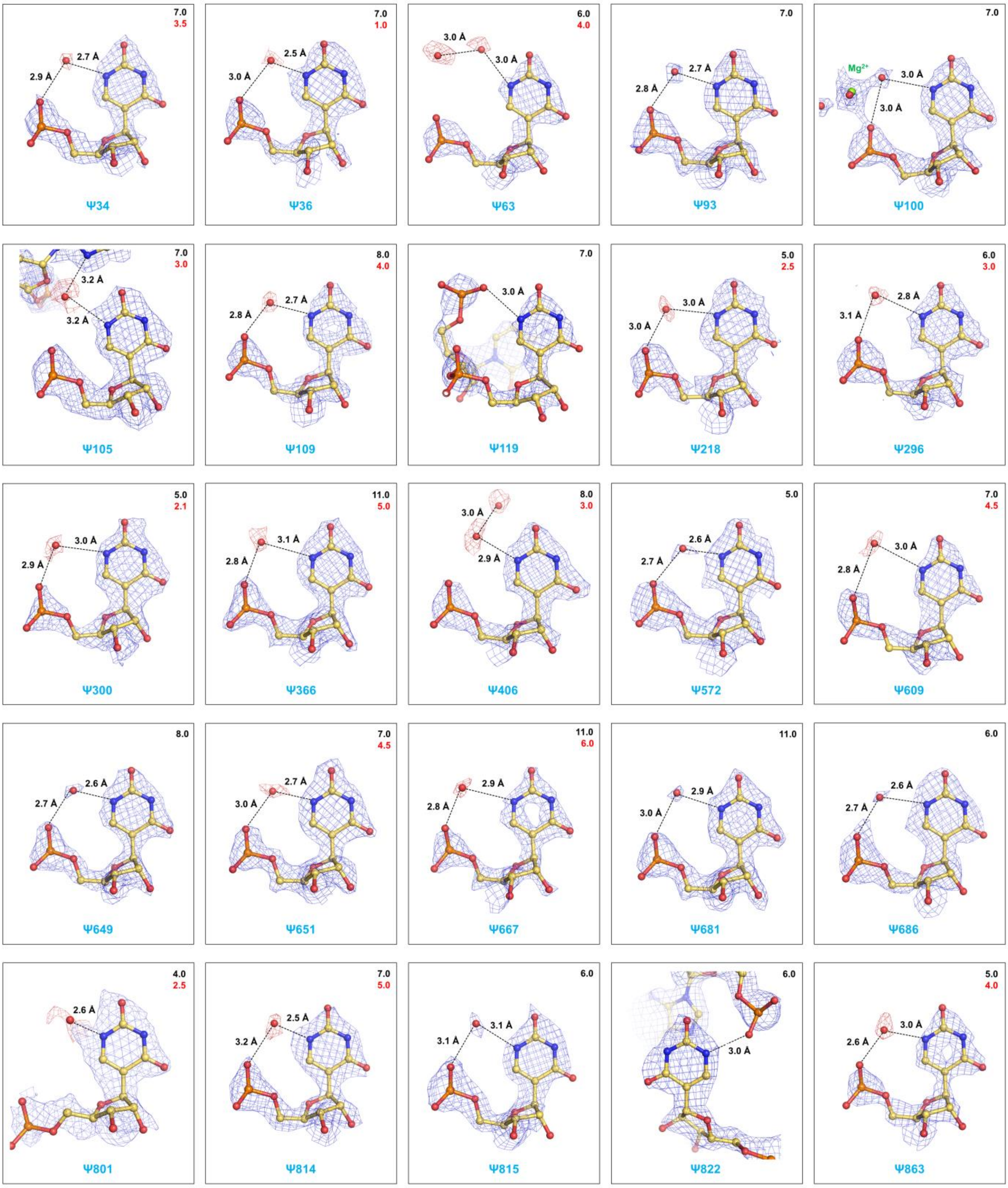

**18S rRNA – pseudo-uridines (cont.)**

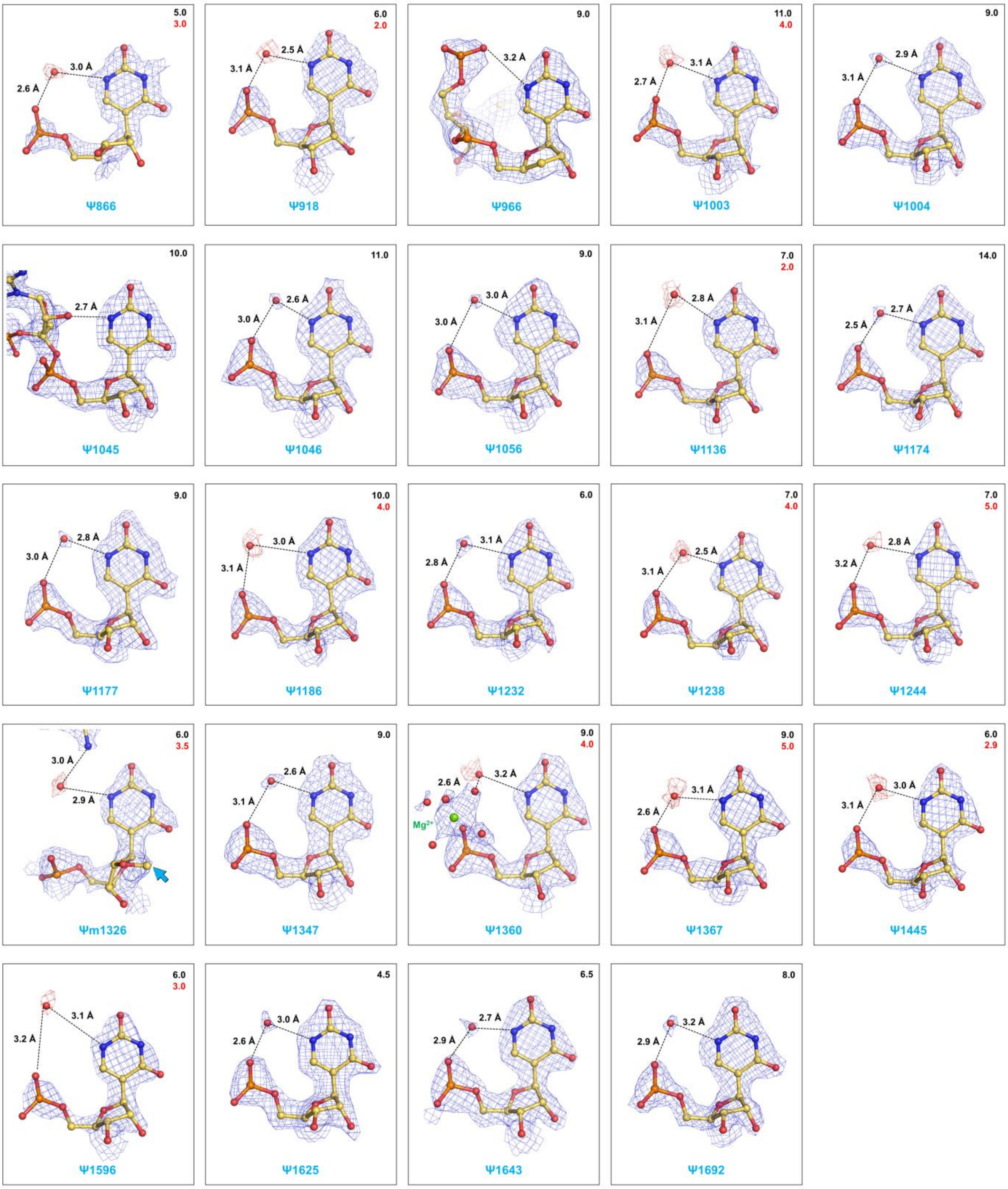

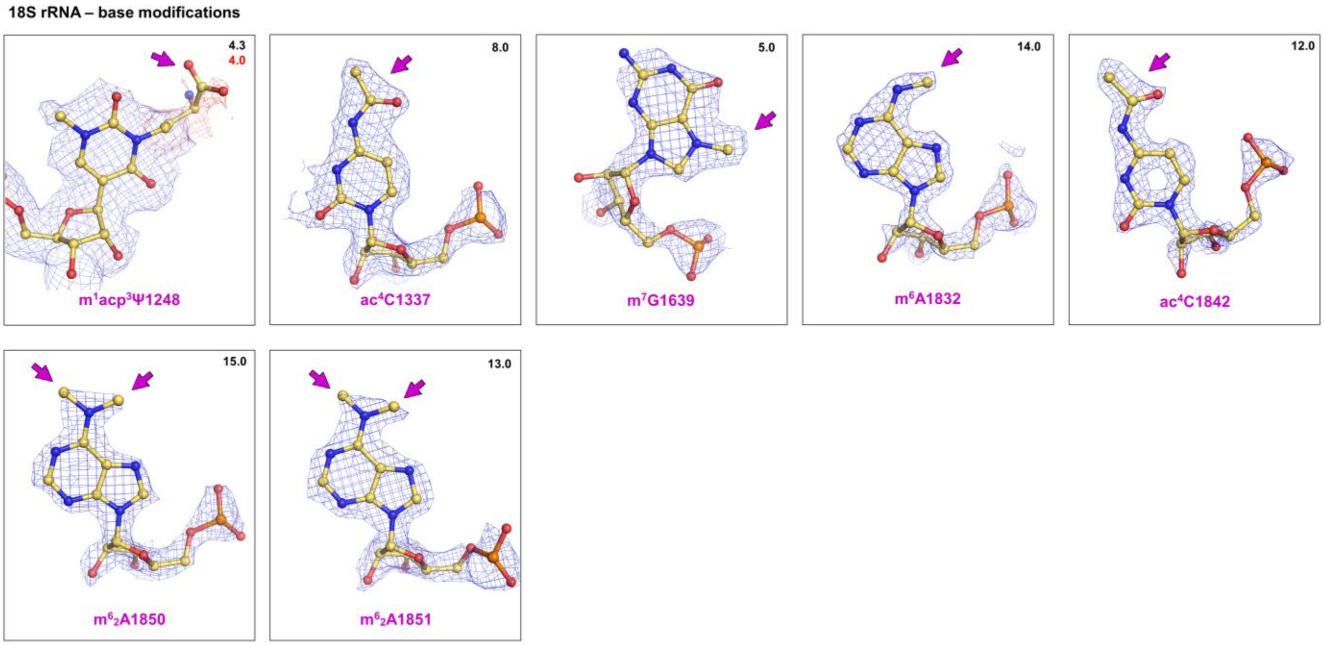

**Chemical modifications seen in the cryo-EM map**

2’-O-Me’s, ψ’s & nucleotide base modifications in the 40S ribosomal subunit (human 18S rRNA)

**Extended Data Fig. 4**

**5.8S rRNA – 2'-O-methylations**

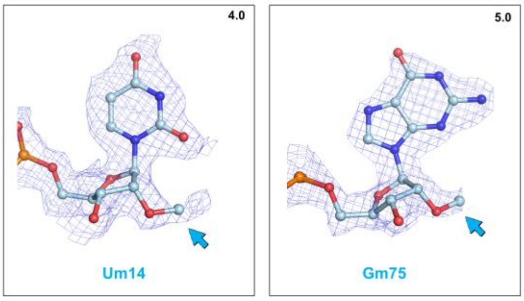

**28S rRNA – 2'-O-methylations**

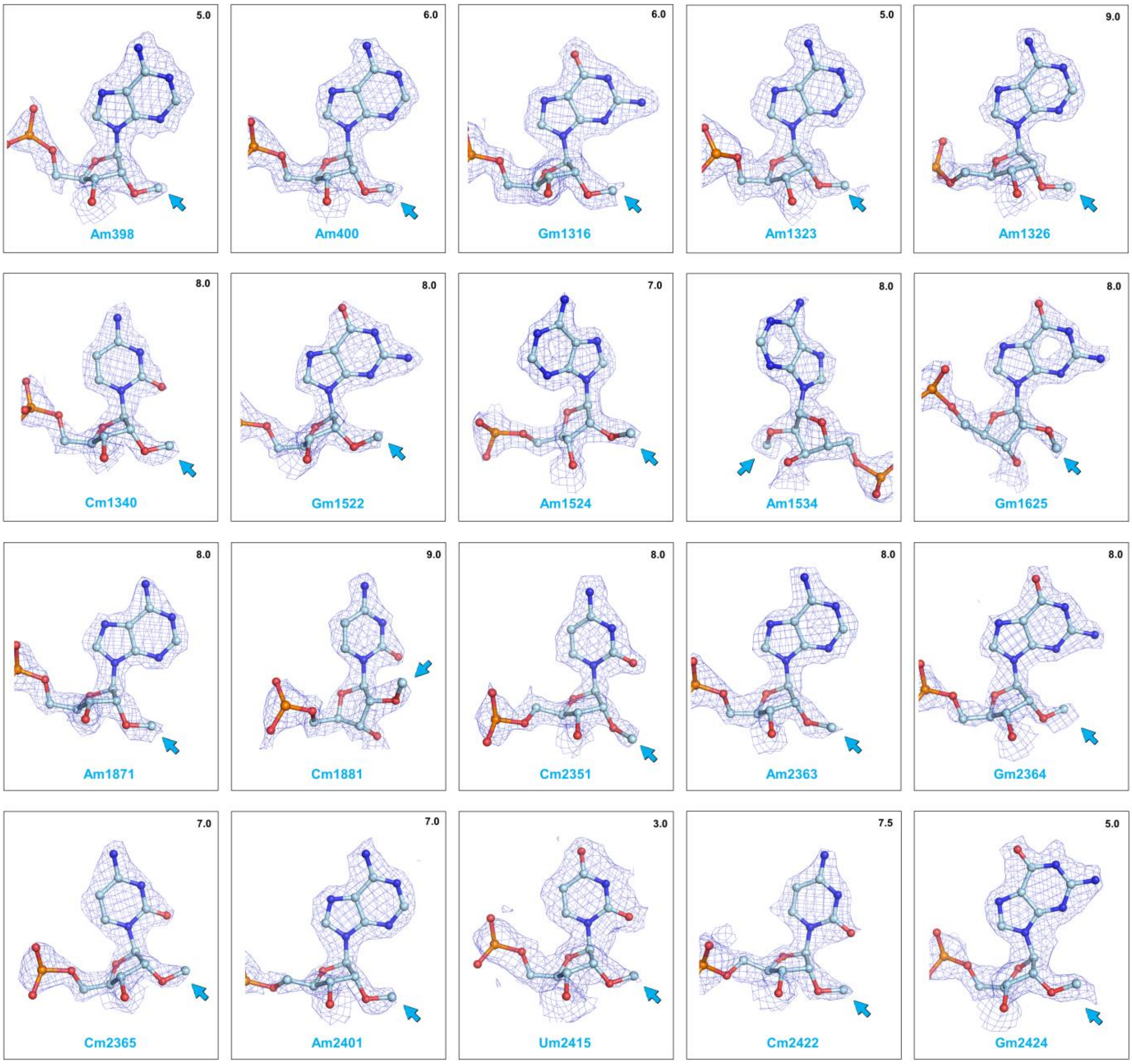

28S rRNA – 2'-O-methylations (cont.)

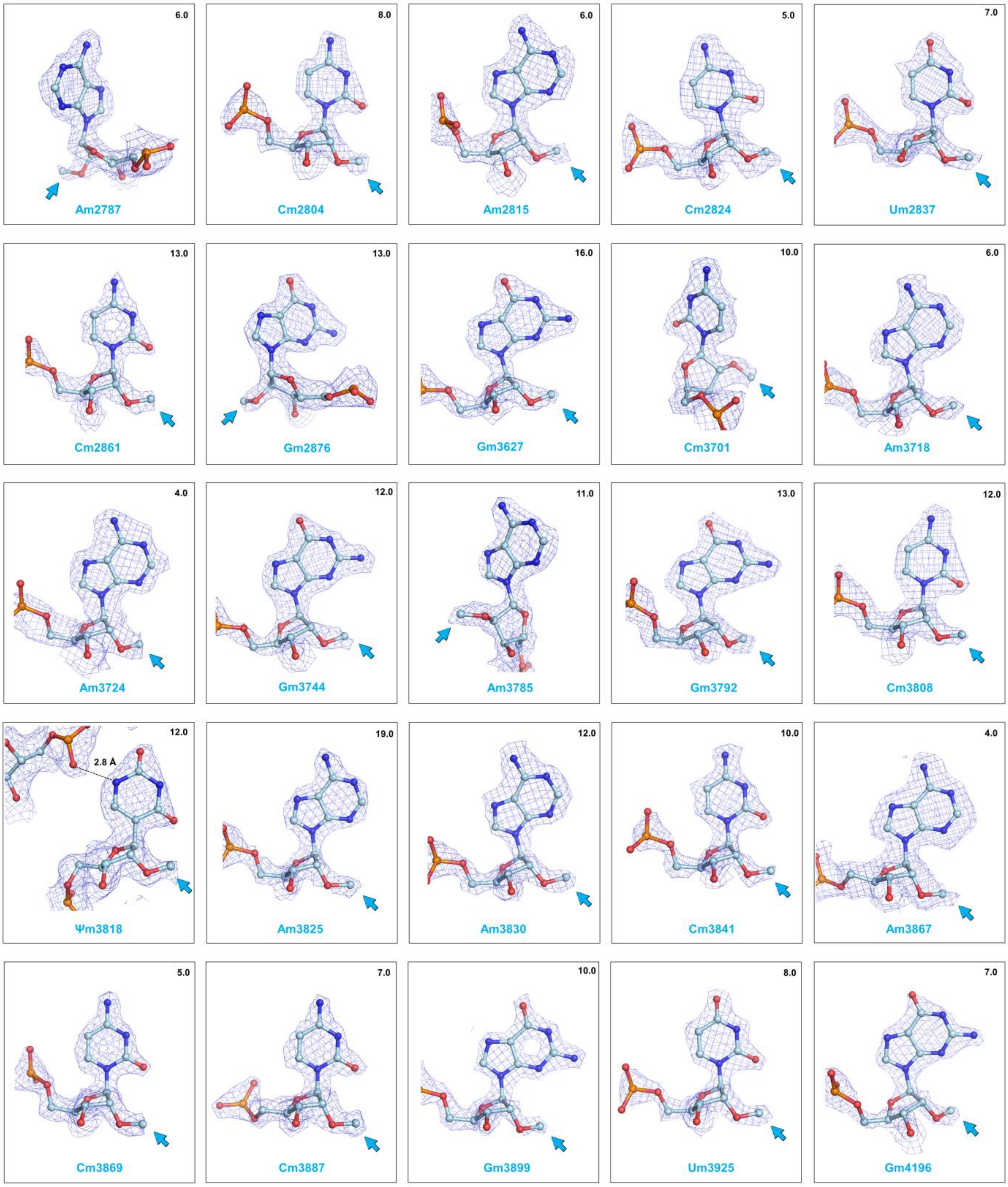

28S rRNA – 2'-O-methylations (cont.)

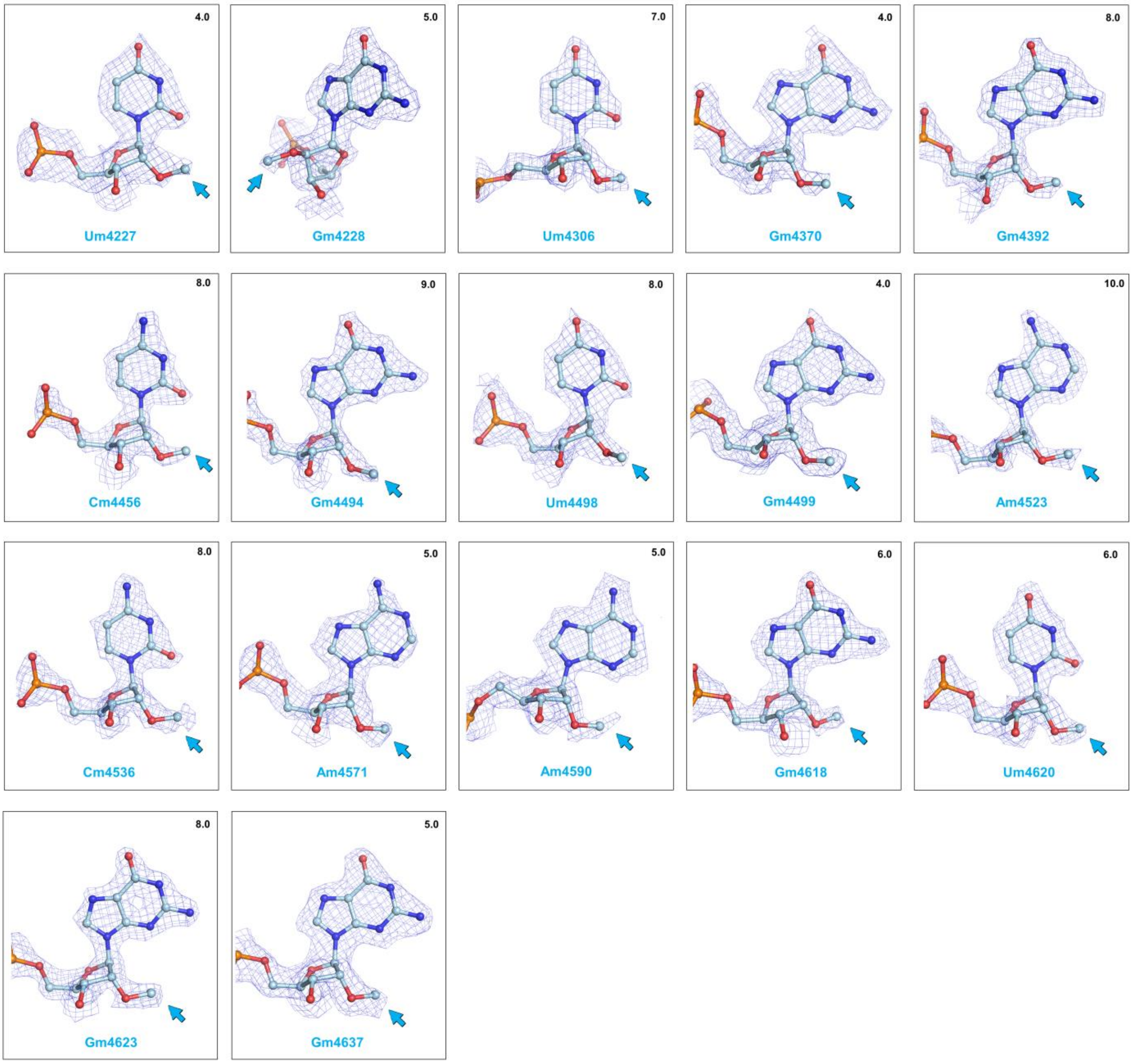

5.8S rRNA – pseudo-uridines

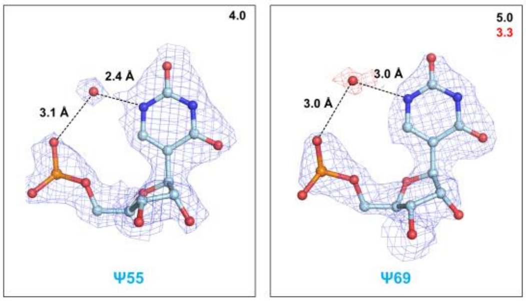

28S rRNA – pseudo-uridines

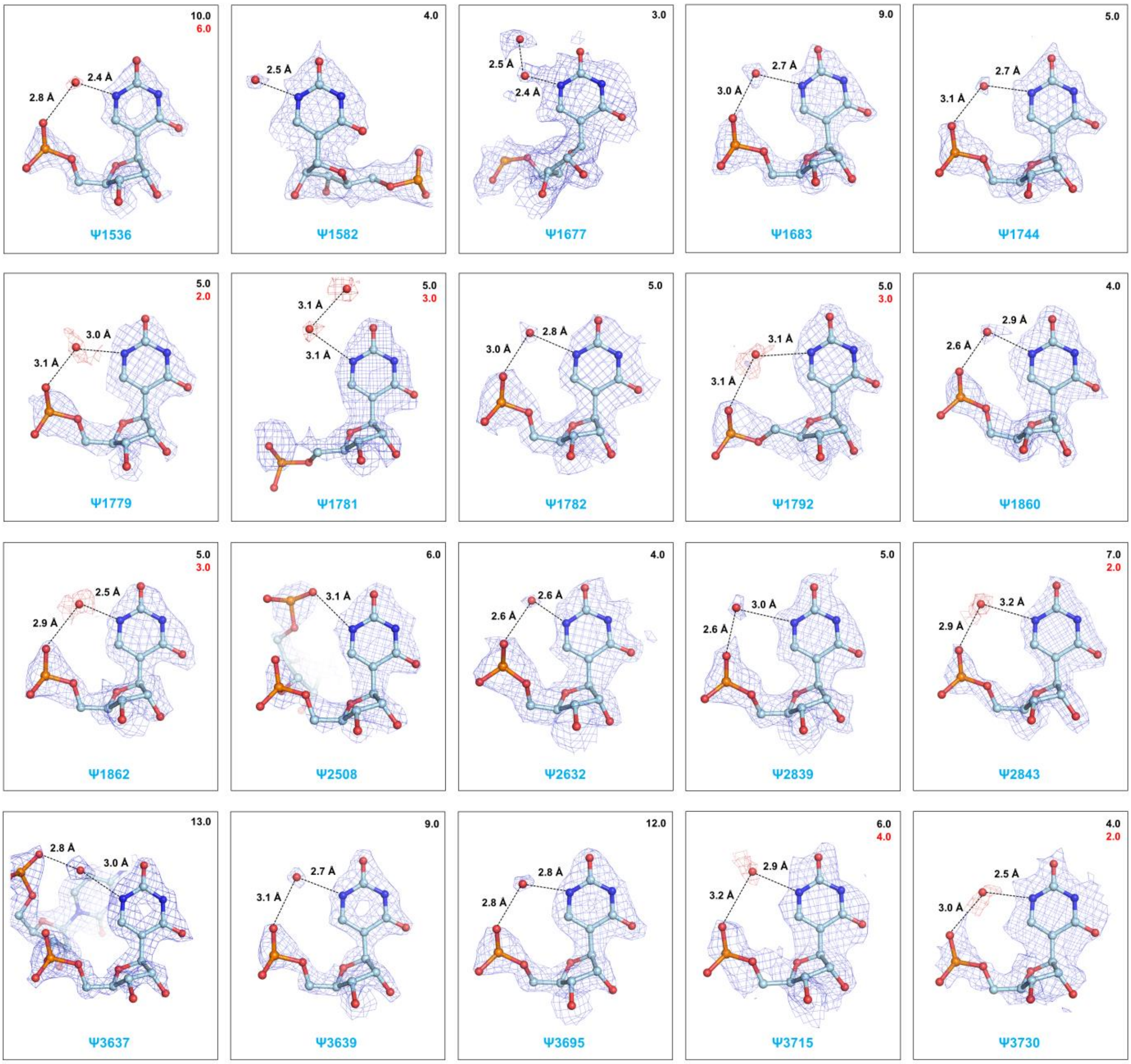

28S rRNA – pseudo-uridines (cont.)

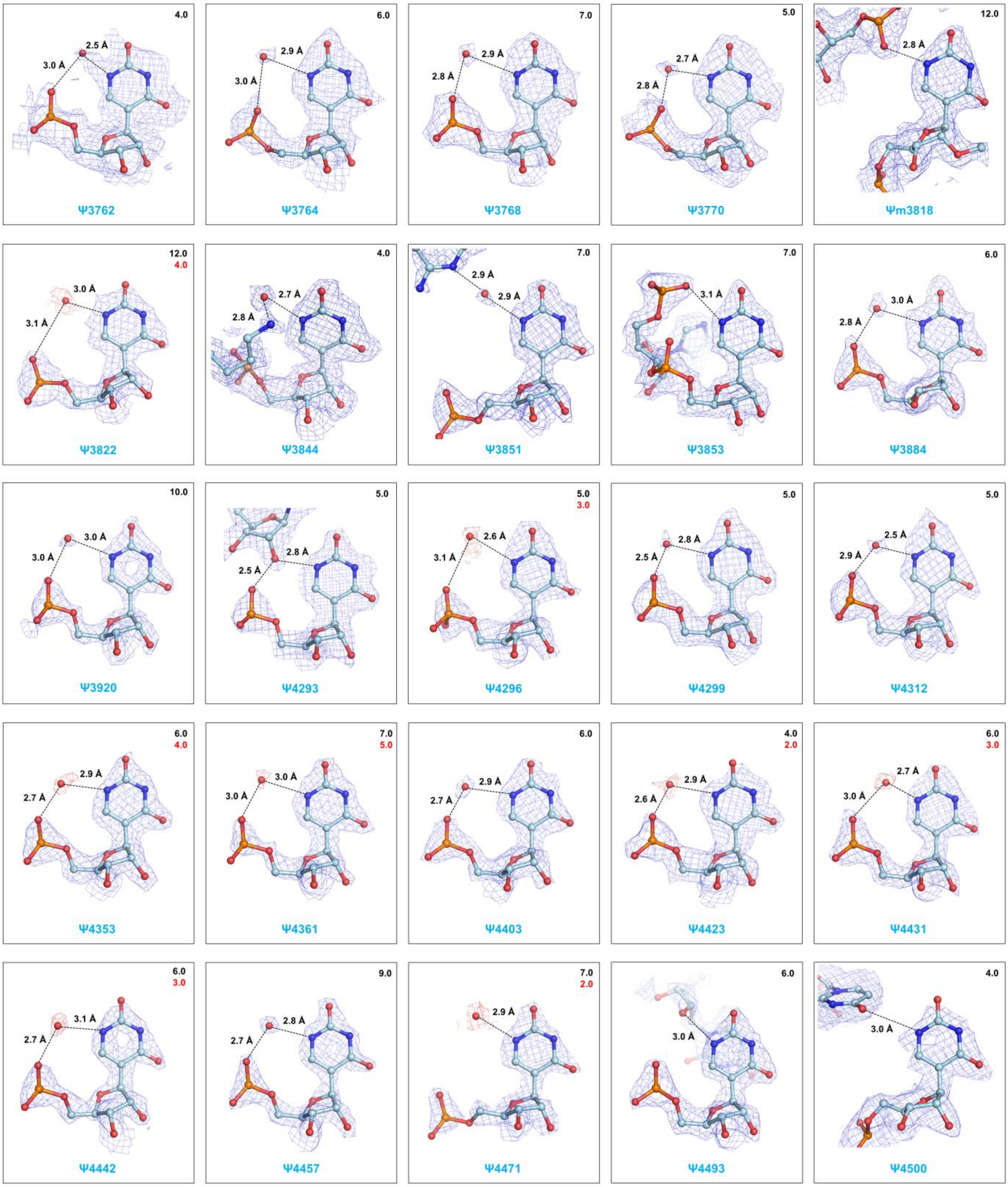

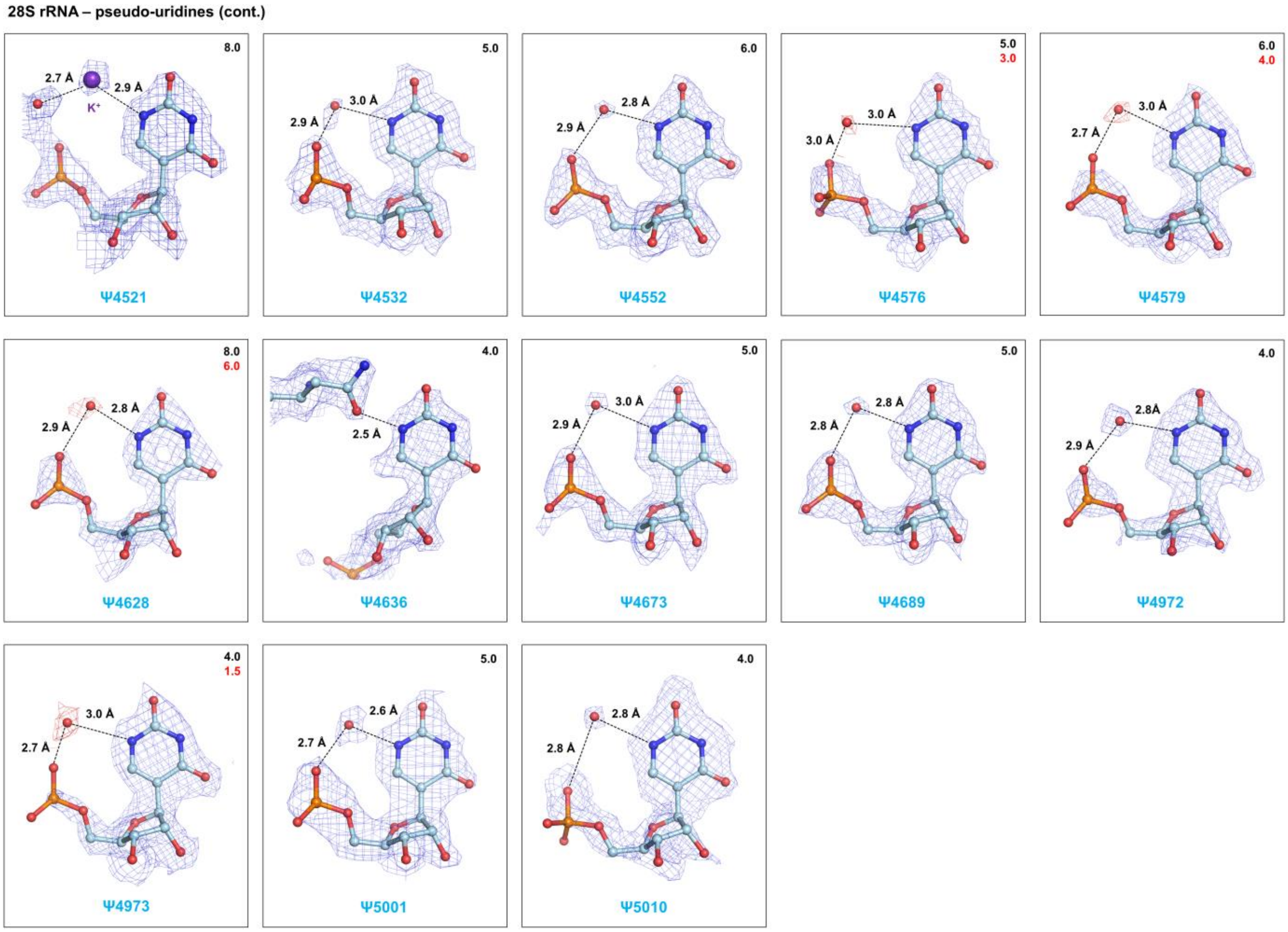

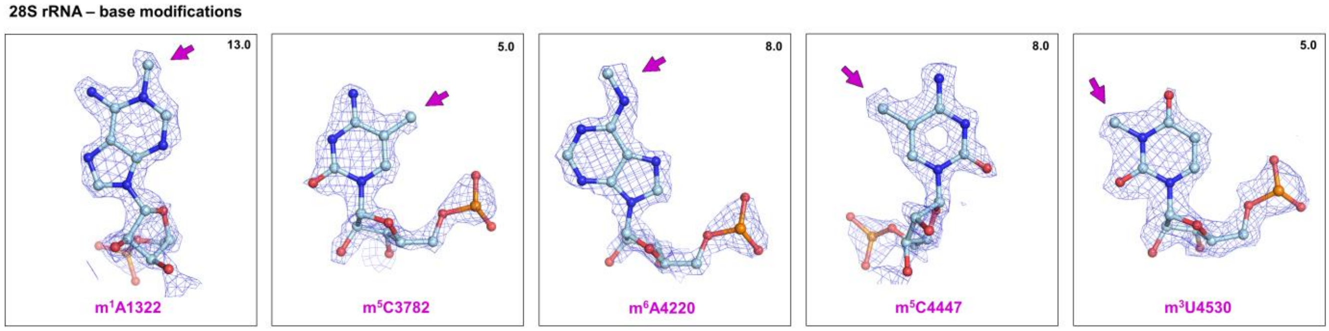

**Chemical modifications seen in the cryo-EM map**

2'-O-Me's, ψ's & nucleotide base modifications in the 60S ribosomal subunit (human 5.8S & 28S rRNA)

**Extended Data Fig. 5**

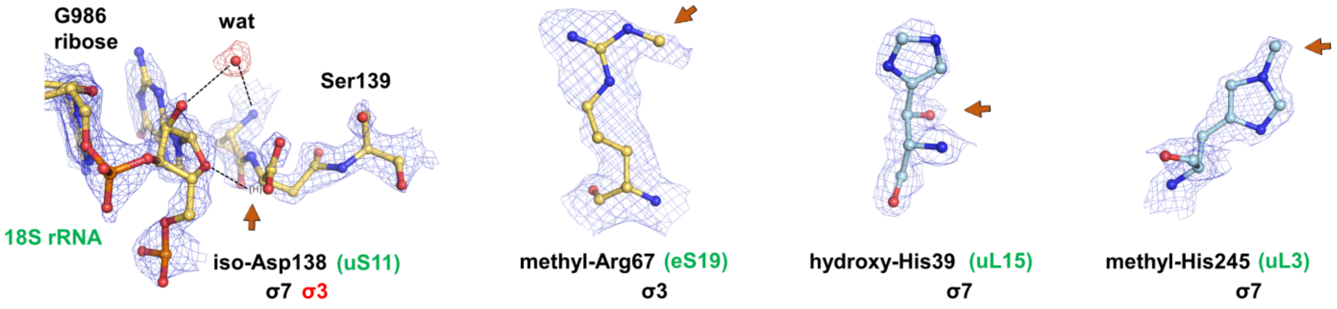

**Post-translational modifications of ribosomal proteins in the human ribosome**

Iso-Asp138 (uS11), methyl-arginine (ribosomal protein eS19), hydroxy-histidine (uL15) and methyl-histidine (uL3).

**Extended Data Fig. 6**

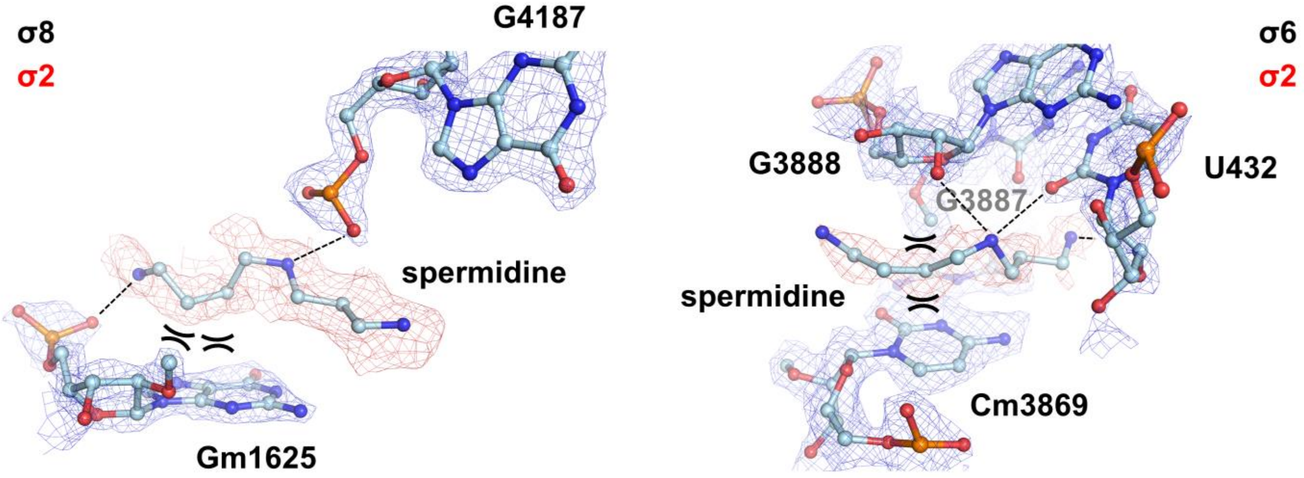

**Visualization of endogenous polyamines (2 spermidines) in the human ribosome**

**Extended Data Fig. 7**

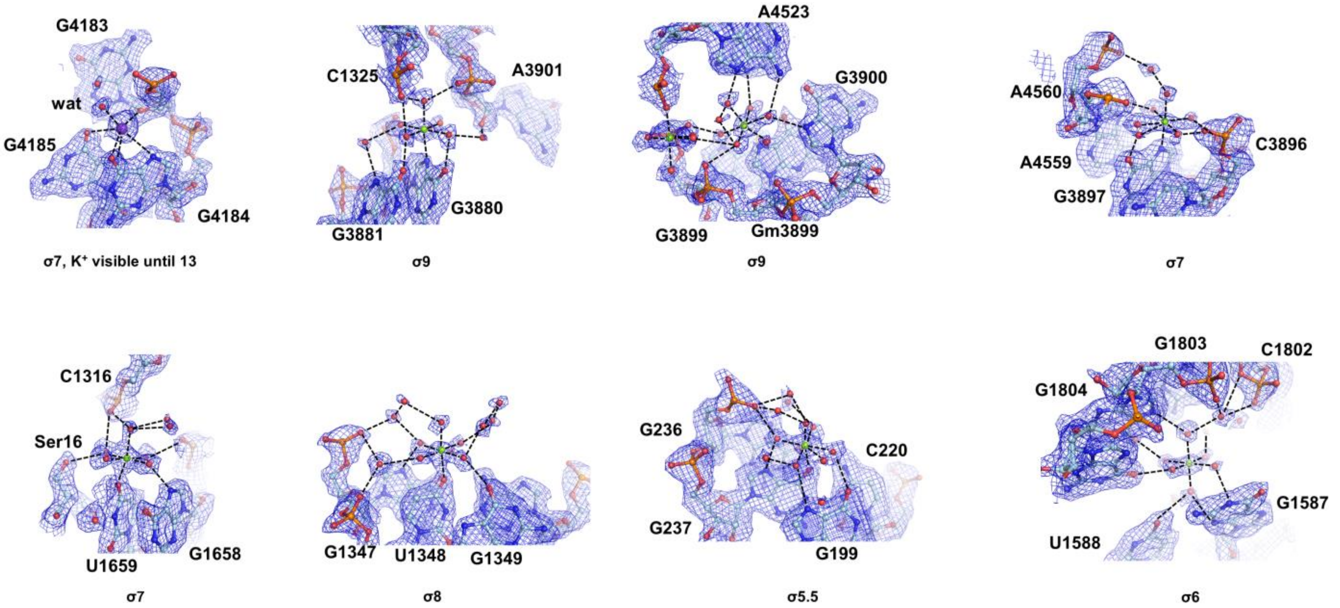

**Assignment of octahedral $Mg^{2+}$ coordinations with water molecules at N7, O6 and O4 positions of nucleotide bases**

Confirmation of $Mg^{2+}$-coordinated rRNA bases that were identified previously by electrostatic potential map calculations[48]; respective σ-levels are indicated.

**Extended Data Fig. 8**

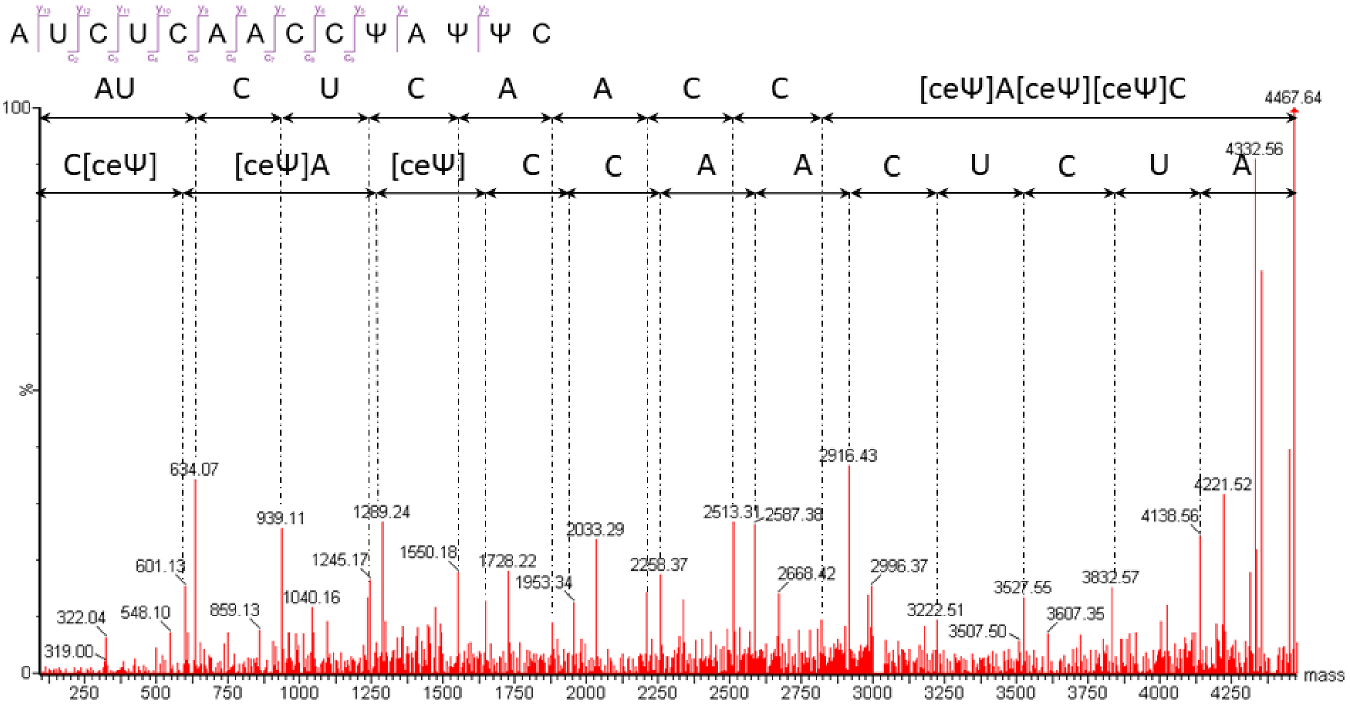

**MS analysis using chimeric oligo nucleotides**

Deconvoluted MS/MS spectrum of [1770]AUCUCAACC[Ψ]A[Ψ][Ψ][1783]C from HeLa 28S rRNA with fragmentation assignments

**Extended Data Fig. 9**

| rRNA | Oligonucleotide name | Sequence (5'-->3') | GC-content |
|---|---|---|---|
| 18S | 18S 253 | CCGCCCCCCGGCCGGGGCCG | 100% |
| 18S | 18S 253-528 | GCCTCGAAAGAGTCCTGTAT | 50% |
| 18S | 18S 528-727 | AGGCAAGGGGCGGGGACGGG | 80% |
| 18S | 18S 727-922 | CGAATGCCCCCGGCCGTCCC | 80% |
| 18S | 18S 922-1187 | CCGTCAATTCCTTTAAGTTT | 35% |
| 18S | 18S 1187-1661 | GTTACCCGCGCCTGCCGGCG | 80% |
| 28S | 28S 249 | CCCGGGCCCGGCGCGCCGGG | 100% |
| 28S | 28S 249-505 | CGGGGGGCGGGAAAGATCCG | 75% |
| 28S | 28S 505-792 | CCGGGGGGCGGAGACGGGGG | 90% |
| 28S | 28S 792-1028 | CGCGCCCCCGCGGGGGAGAC | 90% |
| 28S | 28S 1028-1279 | CGACGTCGCCGCCGACCCCG | 85% |
| 28S | 28S 1279-1538 | TGGCTTCGCCCTGCCCAGGC | 75% |
| 28S | 28S hyb 1538-1772 | [Am][Am][Am][Gm][Um][Um][Um][Gm]<br>[Am](GAAT)[Am][Gm][Gm][Um][Um][Gm][Am] | 30% |
| 28S | 28S 1538-1815 | CACGCCCGGCTCCACGCCAG | 80% |
| 28S | 28S 1815-2061 | GACGGCCGGGTATGGGCCCG | 80% |
| 28S | 28S 2061-2329 | TCTGCACCTGCGGCGGCTCC | 75% |
| 28S | 28S 2329-2554 | CTGGACGCCTCGCGGCGCCC | 85% |
| 28S | 28S 2554-2837 | TGCCGACTTCCCTTACCTAC | 55% |
| 28S | 28S 2554-2881 | CAGCCCTTAGAGCCAATCCT | 55% |
| 28S | 28S 2837-3107 | CCCCCCGCCGCCGCCCCCAC | 95% |
| 28S | 28S 3107-3402 | CCCCGCCCCGCCGCCCGCCG | 100% |
| 28S | 28S 3402-3663 | GTCAACACCCGCCGCGGGCC | 80% |
| 28S | 28S 3663-3915 | GCCAGACTAGAGTCAAGCTC | 55% |
| 28S | 28S 3663-3959 | GGGGGCCTCCCACTTATTCT | 60% |
| 28S | 28S 3915-4157 | GGCACTGTCCCCGGAGCGGG | 80% |
| 28S | 28S 4157-4416 | AAGCGACGTCGCTATGAACG | 55% |
| 28S | 28S 4157-4500 | CGTTCCCTATTAGTGGGTGA | 50% |
| 28S | 28S 4391-4643 | CCGCCACAAGCCAGTTATCC | 60% |
| 28S | 28S 4416-4643 | CGCCCCATTGGCTCCTCAGC | 70% |
| 28S | 28S 4643-4876 | CGTTTCCCAGGACGAAGGGC | 65% |

**RNA & DNA oligos used for mass spectrometry analysis**

Deoxyribonucleotides are in parenthesis; Nm corresponds to 2'-O-methylated nucleotides.

**Extended Data Fig. 10**

MS analysis of 2’-O-Me and ψ modifications in the 18S rRNA

| 18S | | | | | | | | |
|---|---|---|---|---|---|---|---|---|
| | RNase $T_1$ | | RNase A | | RNase $U_2$ | | RNase MC1 | |
| modification | Sequenced fragment | measured *m/z* (z) | Sequenced fragment | measured *m/z* (z) | Sequenced fragment | measured *m/z* (z) | Sequenced fragment | measured *m/z* (z) |
| Am27 | CAU[Am]UGp | 975.13 (-2) | | | | | | |
| ψ34 | [ceY]C[ceY]CAAAGp | 1338.19 (-2) | | | | | | |
| ψ36 | [ceY]C[ceY]CAAAGp | 1338.19 (-2) | | | | | | |
| ψ63 | | | AAG[ceY]ACp | 1006.14 (-2) | | | | |
| Am99 | CUC[Am][ceY]UAAAUCAGp | 1403.87 (-3) | | | CUC[Am][ceY]UA>p | 1125.64 (-2) | | |
| ψ100 | CUC[Am][ceY]UAAAUCAGp | 1403.87 (-3) | | | CUC[Am][ceY]UA>p | 1125.64 (-2) | | |
| ψ105 | CUC[Am]UUAAA[ceY]CAGp | 1403.85 (-3) | | | | | | |
| ψ109 | [ceY]UAUGp | 830.59 (-2) | | | | | | |
| Um116 | U[Um]CC[ceY]U[Um]Gp | 1291.15 (-2) | | | | | | |
| ψ119 | U[Um]CC[ceY]U[Um]Gp | 1291.15 (-2) | | | | | | |
| Um121 | U[Um]CC[ceY]U[Um]Gp | 1291.15 (-2) | | | | | | |
| Am159 | UA[Am]UUCUAGp | 1445.67 (-2) | | | [Am]UUCUA>p | 946.6 (-2) | | |
| Am166 | AG[Am]GCp | 841.62 (-2) | | | | | | |
| Um172 | CUAA[Um]A[Cm]AUGp | 1077.47 (-3) | AA[Um]A[Cm]AUp | 1138.7 (-2) | | | | |
| Cm174 | CUAA[Um]A[Cm]AUGp | 1077.47 (-3) | AA[Um]A[Cm]AUp | 1138.7 (-2) | | | | |
| ψ210 | | | GGGGGGGA[ceY]p | 1039.51 (-3) | | | | |
| ψ218 | CA[ceY]UUAUCAGp | 1078.16 (-3) | | | | | | |
| ψ296 | ACUC[ceY]AGp | 1147.22 (-2) | | | | | | |
| ψ300 | A[ceY]AACCUCGp | 1464.29 (-2) | | | | | | |
| ψ366 | CCCUAUCAAC[ceY]UUCGp | 1586.96 (-3) | | | | | | |
| Um428 | AU[Um]CCGp | 963.12 (-2) | | | | | | |
| Gm436 | GGAGA[Gm]GGAGCp | 1245.55 (-3) | | | | | | |
| Cm462 | CUACCACAU[Cm]CAAGp | 1487.21 (-3) | | | | | | |
| Am468 | | | AAGG[Am]AGGCp | 1515.79 (-2) | | | | |
| Am484 | C[Am]AAUUACCCACUCCCG>p | 1339.68 (-4) | | | | | | |
| Gm509 | | | GGGGA[Gm]GUp | 1367.75 (-2) | | | | |
| Am512 | [Am]G[ceY]GA[Cm]GAAAAAUp | 1450.54 (-3) | | | | | | |
| ψ514 | [Am]G[ceY]GA[Cm]GAAAAAUp | 1450.54 (-3) | | | | | | |
| Cm517 | [Am]G[ceY]GA[Cm]GAAAAAUp | 1450.54 (-3) | | | | | | |
| ψ556 | UAA[ceY]UGp | 995.12 (-2) | | | | | | |
| Am590 | [Am]UCCAUUGp | 1280.66 (-2) | GAGG[Am]Up | 1014.68 (-2) | [Am]UCCA>p | 793.09 (-2) | | |
| Gm601 | | | GGAG[Gm]GCp | 1194.73 (-2) | | | | |
| Um627 | [Um]AAUUCCAGp | 1445.19 (-2) | GG[Um]AAUp | 995.17 (-2) | | | | |
| Gm644 | CUCCAAUA[Gm]CGp | 1179.49 (-3) | | | | | | |
| ψ649 | UA[ceY]A[ceY]UAAAGp | 1111.81 (-3) | | | | | | |
| ψ651 | UA[ceY]A[ceY]UAAAGp | 1111.81 (-3) | | | | | | |
| ψ667 | U[ceY][Am]AAAAGp | 1342.7 (-2) | [ceY][Am]AAAAGCp | 1342.25 (-2) | | | | |
| Am668 | U[ceY][Am]AAAAGp | 1342.7 (-2) | [ceY][Am]AAAAGCp | 1342.25 (-2) | | | U[Am]AAAAGC>p | 1306.68 (-2) |
| ψ686 | A[ceY]CUUGp | 983.11 (-2) | | | | | | |
| ψ769/770[a] | CUC[ceY]UAGp or CUCU[ceY]AGp | 1135.7 (-2) | | | | | | |
| Cm797 | | | GAAG[Cm]G[Um]Up | 1327.23 (-2) | | | | |
| Um799 | [Um]U[ceY]AC[ceY]UUGp | 1475.76 (-2) | GAAG[Cm]G[Um]Up | 1327.23 (-2) | | | | |
| ψ801 | [Um]U[ceY]AC[ceY]UUGp | 1475.76 (-2) | | | | | | |
| ψ804 | [Um]U[ceY]AC[ceY]UUGp | 1475.76 (-2) | | | | | | |
| ψ814 | AAAAAA[ceY][ceY]AGp | 1127.22 (-3) | GAAAAAA[ceY]p | 1347.26 (-2) | | | | |
| ψ815 | AAAAAA[ceY][ceY]AGp | 1127.22 (-3) | | | | | | |
| ψ822 | UG[ceY]UCAAAGp | 1484.78 (-2) | | | | | | |
| ψ863 | AA[ceY]AA[ceY][Gm]Gp | 1377.29 (-2) | AGGAA[ceY]p | 1026.19 (-2) | | | | |
| ψ866 | AA[ceY]AA[ceY][Gm]Gp | 1377.29 (-2) | | | | | | |
| Gm867 | AA[ceY]AA[ceY][Gm]Gp | 1377.29 (-2) | | | | | | |
| ψ918 | AU[ceY]AAGp | 1006.68 (-2) | | | | | | |
| ψ966 | AAAU[ceY]CUUGp | 1465.28 (-2) | | | | | | |
| ψ1002/**1003**[a] | CAU[ceY][ceY]Gp | 1009.68 (-2) | | | | | | |
| ψ1004 | CAU[ceY][ceY]Gp | 1009.68 (-2) | | | | | | |
| Am1031 | | | AAGA[Am]Cp | 998.21 (-2) | | | | |
| ψ1045 | | | GGAGG[ceY]p | 1042.18 (-2) | | | | |
| ψ1061 | A[ceY]ACCGp | 994.18 (-2) | | | | | | |
| ψ1081 | ACCA[ceY]AAACGp | 1093.21 (-3) | | | | | | |
| ψ1186 | | | [ceY]GAAACp | 1006.2 (-2) | | | | |
| Um1288 | | | AGGA[Um]Up | 995.16 (-2) | | | | |
| ψm1326[b] | | | GG[Um]G[Gm]Up | 1018.17 (-2) | | | | |
| Gm1328 | | | GG[Um]G[Gm]Up | 1018.17 (-2) | | | | |
| ψ1358/1359/**1360**[a] | AUU[ceY]Gp | 830.63 (-2) | | | | | | |
| Am1383 | | | | | | | UAACGA[Am]CGAGAC>p | 1413.83 (-3) |
| Cm1391 | ACU[Cm]UGp | 963.17 (-2) | | | CU[Cm]UGG>p | 962.11 (-2) | | |
| Um1442 | | | | | CU[Um]CU[ceY]A>p | 1114.13 (-2) | | |
| ψ1445 | | | | | CU[Um]CU[ceY]A>p | 1114.13 (-2) | | |
| Gm1447 | | | A[Gm]AGGGACp | 1351.27 (-2) | | | | |
| Gm1490 | CAAUAACA[Gm]Gp | 1093.48 (-3) | | | | | | |
| ψ1596 | AACCCCAU[ceY]CGp | 1179.22 (-3) | | | | | | |
| ψ1625 | CAAUUAU[ceY]CCCCAUGp | 1594.87 (-3) | | | U[ceY]CCCCA>p | 1106.14 (-2) | | |
| ψ1643 | [m$^7$G]AAU[ceY]CCCAG>p | 1089.47 (-3) | | | | | | |
| Am1678 | AUU[Am]AGp | 987.14 (-2) | | | | | U[Am]AGUCCC>p | 1271.16 (-2) |
| Cm1703 | | | | | CCG[Cm]CCGUCGCUA>p | 1374.17 (-3) | | |
| Um1804 | AACU[Um]Gp | 975.12 (-2) | | | | | | |
| ψ1839 | | | | | U[Y]UC[ac4C]G>p | 983.1 (-3) | | |

[a]The position of the cyanoethylation could not be determined by fragmentation spectra, but the residues are **resolved** in the structure: ψ1358/1359/**1360**; ψ1002/**1003**.

[b]The cyanotethylated precursor ion was absent from the MS analysis suggesting a Um instead of ψm.

### MS analysis of 2'-O-Me and ψ modifications in the 28S rRNA

| 28S | | | | | | | | |
|---|---|---|---|---|---|---|---|---|
| | RNase $T_1$ | | RNase A | | RNase $U_2$ | | RNase MC1 | |
| modification | Sequenced fragment | measured *m/z* (z) | Sequenced fragment | measured *m/z* (z) | Sequenced fragment | measured *m/z* (z) | Sequenced fragment | measured *m/z* (z) |
| Am398 | | | GAAG[Am]G[Am]GAGUp | 1239.84 (-3) | | | | |
| Am400 | | | GAAG[Am]G[Am]GAGUp | 1239.84 (-3) | | | | |
| Gm1316 | ACCC[Gm]UCUUGp | 1062.13 (-3) | | | CCC[Gm]UCUUG>p | 1420.15 (-2) | | |
| Am1323 | [m$^1$A][Am]AC[Am]CGp | 1164.68 (-2) | | | | | | |
| Am1326 | [m$^1$A][Am]AC[Am]CGp | 1164.68 (-2) | | | | | | |
| Cm1340 | U[Cm]UAACACGp | 1444.69 (-2) | | | | | | |
| Gm1522 | ACCC[Gm]A[Am]AGp | 1482.72 (-2) | [Gm]A[Am]AGAUp | 1178.17 (-2) | CCC[Gm]Ap | 809.61 (-2) | UAGGACCC[Gm]A[Am]AGA>p | 1533.56 (-3) |
| Am1524 | ACCC[Gm]A[Am]AGp | 1482.72 (-2) | [Gm]A[Am]AGAUp | 1178.17 (-2) | | | UAGGACCC[Gm]A[Am]AGA>p | 1533.56 (-3) |
| Am1534 | A[Am]C[ceY]AUGp | 1166.17 (-2) | | | [Am]C[ceY]Ap | 676.10 (-2) | | |
| ψ1536 | A[Am]C[ceY]AUGp | 1166.17 (-2) | | | [Am]C[ceY]Ap | 676.10 (-2) | | |
| ψ1677 | UUUCCC[ceY]CAGp | 1062.11 (-3) | | | | | | |
| ψ1683 | | | AGGA[ceY]p | 861.61 (-2) | | | | |
| ψ1779 | AUCUCAACC[ceY]A[ceY][ceY]C | 1488.51 (-3) | | | | | | |
| ψ1781 | AUCUCAACC[ceY]A[ceY][ceY]C | 1488.51 (-3) | | | | | | |
| ψ1782 | AUCUCAACC[ceY]A[ceY][ceY]C | 1488.51 (-3) | | | | | | |
| ψ1860 | CCAC[ceY]U[ceY]UGp | 978.11 (-3) | | | | | | |
| ψ1862 | CCAC[ceY]U[ceY]UGp | 978.11 (-3) | | | | | | |
| Am1871 | | | [Am]GAACp | 833.63 (-2) | | | | |
| Gm 1946 | | | AGAAAAG[Gm]Up | 1508.23 (-2) | | | | |
| Cm2351 | AU[Cm]UUGp | 963.6 (-2) | | | | | | |
| Am2363 | UA[Am][Gm][Cm]AAUAUUCAAACG>p | 1374.93 (-4) | [Am][Gm][Cm]AAAUp | 1165.18 (-2) | U[Am][Gm][Cm]Ap | 836.13 (-2) | U[Am][Gm][Cm]AAA>p | 1156.17 (-2) |
| Gm2364 | UA[Am][Gm][Cm]AAUAUUCAAACG>p | 1374.93 (-4) | [Am][Gm][Cm]AAAUp | 1165.18 (-2) | U[Am][Gm][Cm]Ap | 836.13 (-2) | U[Am][Gm][Cm]AAA>p | 1156.17 (-2) |
| Cm2365 | UA[Am][Gm][Cm]AAUAUUCAAACG>p | 1374.93 (-4) | [Am][Gm][Cm]AAAUp | 1165.18 (-2) | U[Am][Gm][Cm]Ap | 836.13 (-2) | U[Am][Gm][Cm]AAA>p | 1156.17 (-2) |
| Am2401 | | | GG[Am]GAAGGGUp | 1130.82 (-3) | | | | |
| Um2415 | | | G[Um]GAACp | 994.64 (-2) | | | UG[Um]GAACAG[Cm]A[Gm]U>p | 1421.2 (-3) |
| Cm2422 | [Cm]A[Gm]UUGp | 990.12 (-2) | AG[Cm]A[Gm]Up | 1001.64 (-2) | | | UG[Um]GAACAG[Cm]A[Gm]U>p | 1421.2 (-3) |
| Gm2424 | [Cm]A[Gm]UUGp | 990.12 (-2) | AG[Cm]A[Gm]Up | 1001.64 (-2) | | | UG[Um]GAACAG[Cm]A[Gm]U>p | 1421.2 (-3) |
| ψ2632 | UUCUCUU[ceY]UCUUUGp | 1462.81 (-3) | | | | | | |
| Am2787 | UACCC[Am]UAUCCGp | 1268.15 (-3) | | | | | | |
| Cm2804 | UCU[Cm]CAAGp | 1280.16 (-2) | | | | | | |
| Am2815 | AAC[Am]Gp | 833.63 (-2) | | | | | | |
| Cm2824 | | | GG[Cm]AUp | 830.12 (-2) | | | | |
| Um2837 | AACAA[Um]Gp | 1151.16 (-2) | | | | | | |
| Gm2876 | UAACUUC[Gm]Gp | 1453.17 (-2) | | | | | | |
| Gm3627 | | | AGG[Gm]GAAUp | 1351.69 (-2) | | | | |
| ψ3695 | AUU[ceY]CUGp | 1136.13 (-2) | | | | | | |
| Cm3701 | CC[Cm]AGp | 809.61 (-2) | | | | | | |
| ψ3715 | [ceY]CA[Am]AGp | 1013.15 (-2) | | | | | | |
| Am3718 | [ceY]CA[Am]AGp | 1013.15 (-2) | A[Am]AGUp | 834.11 (-2) | | | | |
| Am3724 | | | GA[Am]GAAAUp | 1335.68 (-2) | | | | |
| Gm3744 | [Gm]UAAACGp | 1159.15 (-2) | | | | | | |
| ψ3758 | [ceY]A[Am]C[ceY]AUGp | 1345.68 (-2) | GGGAG[ceY]p | 1042.15 (-2) | | | | |
| Am3760 | [ceY]A[Am]C[ceY]AUGp | 1345.68 (-2) | | | [Am]C[ceY]A[ceY]GA>p | 1183.66 (-2) | | |
| ψ3762 | [ceY]A[Am]C[ceY]AUGp | 1345.68 (-2) | | | [Am]C[ceY]A[ceY]GA>p | 1183.66 (-2) | | |
| ψ3764 | UA[Am]C[ceY]A[ceY]Gp | 1345.68 (-2) | | | [Am]C[ceY]A[ceY]GA>p | 1183.66 (-2) | | |
| ψ3768 | AC[ceY]C[ceY]CUUAAGp | 1197.49 (-3) | | | C[ceY]C[ceY]CUUA>p | 1286.15 (-2) | | |
| ψ3770 | AC[ceY]C[ceY]CUUAAGp | 1197.49 (-3) | | | C[ceY]C[ceY]CUUA>p | 1286.15 (-2) | | |
| Am3785 | C[m$^5$C]AA[Am]UGp | 1146.16 (-2) | | | | | | |
| Gm3792 | CCUC[Gm]UCAUCUAAUUAGp | 1348.92 (-4) | | | CUC[Gm]UCA>p | 1106.65 (-2) | | |
| Cm3808 | | | GA[Cm]GCp | 829.61 (-2) | | | UAGUGA[Cm]GCGCA>p | 1301.83 (-3) |
| ψ3822 | | | GAA[ceYm]GGAUp / GAA[Um]GGA[ceY]p | 1358.68 (-2) | | | | |
| Am3830 | [Am]UUCCCACUGp | 1056.8 (-3) | GAG[Am]Up | 842.11 (-2) | [Am]UUCCCA>p | 1098.63 (-2) | | |
| Cm3841 | U[Cm]CC[ceY]ACCUACUA[ceY]CCAG>p | 1436.92 (-4) | | | | | | |
| ψ3844 | U[Cm]CC[ceY]ACCUACUA[ceY]CCAG>p | 1436.92 (-4) | | | | | | |
| ψ3851 | U[Cm]CCUACCUAC[ceY]AUCCAG>p | 1423.69 (-4) | | | | | | |
| ψ3853 | U[Cm]CC[ceY]ACCUACUA[ceY]CCAG>p | 1436.92 (-4) | | | | | | |
| Am3867 | AAACCAC[Am]Gp | 1467.72 (-2) | | | | | | |
| Cm3869 | [Cm]CAAGp | 821.62 (-2) | | | | | | |
| Cm3887 | GG[Cm]GGAAUp | 1339.68 (-2) | | | | | | |
| Gm3899 | GG[Gm]GAAAGAAGACp | 1459.53 (-3) | | | | | | |
| Um3925 | AC[Um]CUAG>p | 1118.63 (-2) | | | | | | |
| Gm3944 | AA[Gm]AGp | 853.63 (-2) | GAA[Gm]AGACp | 1343.21 (-2) | | | | |
| Cm4054 | AAAUACCA[Cm]UACUCUG>p | 1269.66 (-4) | | | | | UACCA[Cm]UAC>p | 1415.66 (-2) |
| Gm4196 | | | G[Gm]GGCp | 857.63 (-2) | | | | |
| Um4227 | [Um][Gm]UCCUAAGp | 973.12 (-3) | AGG[Um][Gm]Up | 1010.13 (-2) | | | | |
| Gm4228 | [Um][Gm]UCCUAAGp | 973.12 (-3) | AGG[Um][Gm]Up | 1010.13 (-2) | | | | |
| ψ4299 | A[ceY]UUUCAGp | 1300.64 (-2) | | | | | | |
| Um4306 | | | AG[Um]ACp | 822.11 (-2) | | | | |
| ψ4312 | AA[ceY]ACAGp | 1170.67 (-2) | | | | | | |
| ψ4361 | UUU[ceY]AAGp | 1148.13 (-2) | | | | | | |
| Gm4370 | | | AGGA[Gm]GUp | 1187.16 (-2) | | | | |
| Gm4392 | | | AG[Gm]GAUp | 1014.64 (-2) | | | UACCACAG[Gm]GA>p | 1194.49 (-3) |
| ψ4420 | U[ceY]CAUUAG | 1147.64 (-2) | | | | | | |
| ψ4423 | UUCA[ceY]UAG | 1147.64 (-2) | | | | | | |
| ψ4442 | A[ceY]CCUU[m$^5$C]Gp | 1295.15 (-2) | | | | | | |
| Cm4456 | [Cm][ceY]CUUCCUAUCAUUG>p | 1578.19 (-3) | | | [Cm][ceY]CUUCCUA>p | 1419.16 (-2) | | |
| ψ4457 | [Cm][ceY]CUUCCUAUCAUUG>p | 1578.19 (-3) | | | [Cm][ceY]CUUCCUA>p | 1419.16 (-2) | | |
| Um4498 | AU[Um][Gm][ceY]UCACCCACUAAUAG>p | 1525.68 (-4) | | | U[Um][Gm][ceY]UCAp | 1150.15 (-2) | | |
| Gm4499 | AU[Um][Gm][ceY]UCACCCACUAAUAG>p | 1525.68 (-4) | | | U[Um][Gm][ceY]UCAp | 1150.15 (-2) | | |
| ψ4500 | AU[Um][Gm][ceY]UCACCCACUAAUAG>p | 1525.68 (-4) | | | U[Um][Gm][ceY]UCAp | 1150.15 (-2) | | |
| ψ4532 | [m$^3$U]U[ceY]AGp | 837.61 (-2) | | | | | | |
| Cm4536 | | | AGA[Cm]Cp | 821.62 (-2) | | | | |
| Am4590 | UA[Am]UCCUG>p | 1271.66 (-2) | | | | | | |
| Gm4618 | [Gm]U[Um]CA[Gm]ACAU[ceY]UG>p | 1418.19 (-3) | | | [Gm]U[Um]CAp | 817.61 (-2) | | |
| Um4620 | [Gm]U[Um]CA[Gm]ACAU[ceY]UG>p | 1418.19 (-3) | | | [Gm]U[Um]CAp | 817.61 (-2) | | |
| Gm4623 | [Gm]U[Um]CA[Gm]ACAU[ceY]UG>p | 1418.19 (-3) | | | | | | |
| ψ4628 | [Gm]U[Um]CA[Gm]ACAU[ceY]UG>p | 1418.19 (-3) | | | | | | |
| ψ4636 | UA[ceY][Gm]UG>p | 1001.12 (-2) | | | | | | |
| Gm4637 | UA[ceY][Gm]UG>p | 1001.12 (-2) | | | | | | |
| ψ4673 | CUACCA[ceY]CUGp | 1069.8 (-3) | | | | | | |
| ψ4972 | CUAAACCA[ceY][ceY]CGp | 1306.84 (-3) | | | | | | |
| ψ4973 | CUAAACCA[ceY][ceY]CGp | 1306.84 (-3) | | | | | | |
| ψ5039 | AUCUA[ceY]UGp | 1300.64 (-2) | | | | | | |

MS analysis of 2’-O-Me and ψ modifications in the 5.8S rRNA

| 5.8S | | | | | | | | |
|---|---|---|---|---|---|---|---|---|
| | RNase $T_1$ | | RNase A | | RNase $U_2$ | | RNase MC1 | |
| modification | Sequenced fragment | measured *m/z* (z) | Sequenced fragment | measured *m/z* (z) | Sequenced fragment | measured *m/z* (z) | Sequenced fragment | measured *m/z* (z) |
| Um14 | | | GG[Um]GGAUp | 1175.69 (-2) | | | | |
| ψ55 | C[ceY]Gp | 1026.14 (-1) | | | | | | |
| ψ69 | | | | | | | | |
| Gm75 | AAUU[Gm]CAGp | 1312.17 (-2) | | | UU[Gm]CA>p | 801.59 (-2) | | |

MS analysis of nucleotide base modifications in the 18S rRNA

| 18S | | | | | | | | |
|---|---|---|---|---|---|---|---|---|
| | RNase $T_1$ | | RNase A | | RNase $U_2$ | | RNase MC1 | |
| modification | Sequenced fragment | measured *m/z* (z) | Sequenced fragment | measured *m/z* (z) | Sequenced fragment | measured *m/z* (z) | Sequenced fragment | measured *m/z* (z) |
| m$^{1}$acp$^{3}$ψ1248 | AC[cem$^{1}$acp$^{3}$Y]CAACACGp | 1123.5 (-3) | | | | | | |
| ac$^{4}$C1337 | | | | | C[ac$^{4}$C]GUUCUUA>p | 1426.69 (-2) | | |
| m$^{7}$G1639 | [m$^{7}$G]AAU[ceY]CCCAG>p | 1089.47 (-3) | GAG[m$^{7}$G]AAUp | 1179.22 (-2) | | | | |
| m$^{6}$A1832 | UA[m$^{6}$A]CAAGp | 1151.18 (-2) | | | | | | |
| ac$^{4}$C1842 | UUUC[ac$^{4}$C]Gp | 965.6 (-2) | | | UUUC[ac$^{4}$C]G>p | 956.59 (-2) | | |
| m$^{6}_{2}$A1850 | [m$^{6}_{2}$A][m$^{6}_{2}$A]CCUGp | 995.67 (-2) | G[m$^{6}_{2}$A][m$^{6}_{2}$A]Cp | 690.14 (-2) | UG[m$^{6}_{2}$A][m$^{6}_{2}$A]CCUGCGGAA>p | 1425.55 (-3) | | |
| m$^{6}_{2}$A1851 | [m$^{6}_{2}$A][m$^{6}_{2}$A]CCUGp | 995.67 (-2) | G[m$^{6}_{2}$A][m$^{6}_{2}$A]Cp | 690.14 (-2) | UG[m$^{6}_{2}$A][m$^{6}_{2}$A]CCUGCGGAA>p | 1425.55 (-3) | | |

MS analysis of nucleotide base modifications in the 28S rRNA

| 28S | | | | | | | |
|---|---|---|---|---|---|---|---|
| | RNase $T_1$ | | RNase A | | RNase $U_2$ | | RNase MC1 |
| m$^{1}$A1322 | [m$^{1}$A][Am]AC[Am]CGp | 1164.68 (-2) | | | | | |
| m$^{5}$C3782 | C[m$^{5}$C]AA[Am]UGp | 1146.16 (-2) | | | | | |
| ψm3818 | AA[ceYm]Gp | 696.1 (-2) | GAA[ceYm]GGAUp / GAA[Um]GGA[ceY]p | 1358.68 (-2) | | | |
| m$^{6}$A4220 | UA[m$^{6}$A]CGp | 822.11 (-2) | | | | | |
| m$^{5}$C4447 | A[ceY]CCUU[m$^{5}$C]Gp | 1295.15 (-2) | | | | | |
| m$^{3}$U4530 | [m$^{3}$U]U[ceY]AGp | 837.61 (-2) | | | | | |

**18S, 28S, 5.8S & 5S rRNA fragments with 2’-O-Me’s, ψ’s & nucleotide base modifications analyzed by mass spectrometry analysis**

**Extended Data Fig. 11**

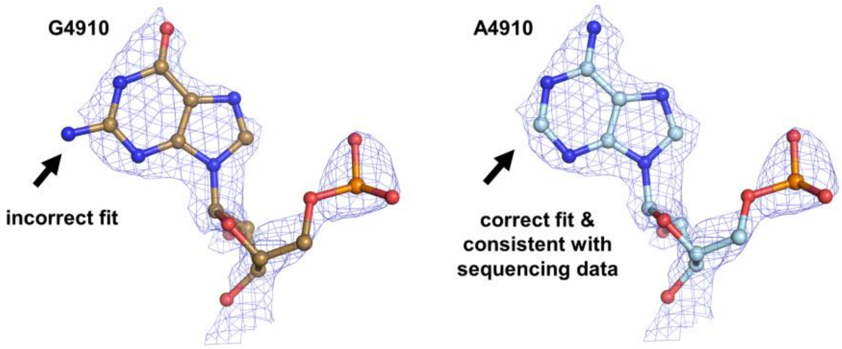

**Correction of the rRNA reference sequence as found by sequencing and confirmed from the structure and MS analysis**

A4910 is labeled in red in the 28S rRNA sequence (see Extended Data Fig. 13).

**Extended Data Fig. 12**

### 18S rRNA sequencing with annotations of chemically modified nucleotides

| | | | | | | | | | | |
|---|---|---|---|---|---|---|---|---|---|---|
| 1 | UACCUGGUUG | AUCCUGCCAG | UAGCAUAmUGC | UUGΨCΨCAAA | GAUUAAGCCA | UGCAUGUCUA | AGΨACGCACG | GCCGGUACAG | UGAAACUGCG | AAΨGGCUCAmΨ |
| 101 | UAAAΨCAGΨU | AUGGUUmCCΨU | UmGGUCGCUCG | CUCCUCUCCU | ACUUGGAUAA | CUGUGGUAAmU | UCUAGAmGCUmA | AUmACmAUGCCG | ACGGGCGCUG | ACCCCCUUCG |
| 201 | CGGGGGGGAΨ | GCGUGCAΨUU | AUCAGAUCAA | AACCAACCCG | GUCAGCCCCU | CUCCGGCCCC | GGCCGGGGGG | CGGGCGCCGG | CGGCUUUGGU | GACUCΨAGAΨ |
| 301 | AACCUCGGGC | CGAUCGCACG | CCCCCCGUGG | CGGCGACGAC | CCAUUCUAAC | GUCUmGCCCUA | UCAACΨUUCG | AUGGUAGUCG | CCGUGCCUAC | CAUGGUGACC |
| 401 | ACGGGΨGACG | GGGAAUCAGG | GUUCGAUUmCC | GGAGAGmGGAG | CCUGAGAAAC | GGCUACCACA | UCmCAAGGAmAG | GCAGCAGGCG | CGCAmAAUUAC | CCACUCCCGA |
| 501 | CCCGGGGAGmG | UAmGΨGACmGAA | AAAUAACAAU | ACAGGACUCU | UUCGAGGCCC | UGUAAΨUGGA | AUGAGUCCAC | UΨUAAAmUCCU | UUAACGAGGAm | UCCAUUGGAG |
| 601 | GmGCAAGUCΨG | GUGCCAGCAG | CmCGCGGUmAAU | UCCAGCUCCA | AUAGmCGUAΨA | ΨUAAAGUUGC | UGCAGUΨAmAA | AAGCUCGUAG | ΨUGmGAΨCUUG | GGAGCGGGCG |
| 701 | GGCGGUCCGC | CGCGAGGCGA | GCCACCGCCC | GUCCCCGCCC | CUUGCCUCUC | GGCGCCCCCU | CGAUGCUCΨΨ | AGCUGAGUGU | CCCGCGGGGC | CCGAAGCmGUmU |
| 801 | ΨACΨUUGAAA | AAAΨΨAGAGU | GΨUCAAAGCA | GGCCCGAGCC | GCCUGGAUAC | CGCAGCUAGG | AAΨAAΨGmGAA | UAGGACCGCG | GUUCUAUUUU | GUUGGUΨUUC |
| 901 | GGAACUGAGG | CCAUGAUΨAA | GAGGGACGGC | CGGGGGCAUU | CCGUAUUGCG | CGCUAGAGGU | GAAAUΨCUUG | GACCGGCGCA | AGACGGACCA | GAGCGAAAGC |
| 1001 | AUΨUGCCAAG | AAUGUUUUCA | UUAAUCAAGA | AmCGAAAGUCG | GAGGΨΨCGAA | GACGAΨCAGA | ΨACCGUCGUA | GUUCCGACCA | ΨAAACGAUGC | CGACCGGCGA |
| 1101 | UGCGGCGGCG | UUAUUCCCAU | GACCCGCCGG | GCAGCΨUCCG | GGAAACCAAA | GUCUUUGGGU | UCCGGGGGGA | GUAΨGGΨUGC | AAAGCΨGAAA | CUUAAAGGAA |
| 1201 | UUGACGGAAG | GGCACCACCA | GGAGUGGAGC | CΨGCGGCΨUA | AUUΨGACm1acp3ΨCA | ACACGGGAAA | CCUCACCCGG | CCmCGGACACG | GACAGGAUmUG | ACAGAUUGAU |
| 1301 | AGCUCUUUCU | CGAUUCCGUG | GGUGGΨmGGmUG | CAUGGCac4CGUU | CUUAGUΨGGU | GGAGCGAUUΨ | GUCUGGΨUAA | UUCCGAUAAC | GAAmCGAGACU | CmUGGCAUGCU |
| 1401 | AACUAGUUAC | GCGACCCCCG | AGCGGUCGGC | GUCCCCCAACm | UUmCUΨAGmAGG | GACAAGUGGC | GUUCAGCCAC | CCGAGAUUGA | GCAAUAACAGm | GUCUGUGAUG |
| 1501 | CCCUUAGAUG | UCCGGGGCUG | CACGCGCGCU | ACACUGACUG | GCUCAGCGUG | UGCCUACCCU | ACGCCGGCAG | GCGCGGGUAA | CCCGUUGAAC | CCCAUΨCGUG |
| 1601 | AUGGGGAUCG | GGGAUUGCAA | UUAUΨCCCCA | UGAACGAGm7GA | AUΨCCCAGUA | AGUGCGGGUC | AUAAGCUUmGC | GUUGAUUAmAG | UCCCUGCCCU | UΨGUACACAC |
| 1701 | CGCmCCGUCGC | UACUACCGAU | UGGAUGGUUU | AGUGAGGCCC | UCGGAUCGGC | CCCGCCGGGG | UCGGCCCACG | GCCCUGGCGG | AGCGCUGAGA | AGACGGUCGA |
| 1801 | ACUUmGACUAU | CUAGAGGAAG | UAAAAGUCGU | Am6ACAAGGUΨU | CCGUAGGUGm6(2)A | m6(2)ACCUGCGGAA | GGAUCAUUA | | | |

### 28S rRNA sequencing with annotations of chemically modified nucleotides

| | | | | | | | | | | |
|---|---|---|---|---|---|---|---|---|---|---|
| 1 | CGCGACCUCA | GAUCAGACGU | GGCGACCCGC | UGAAUUUAAG | CAUAUUAGUC | AGCGGAGGAG | AAGAAACUAA | CCAGGAUUCC | CUCAGUAACG | GCGAGUGAAC |
| 101 | AGGGAAGAGC | CCAGCGCCGA | AUCCCCGCCC | CGCGGCGGGG | CGCGGGACAU | GUGGCGUACG | GAAGACCCGC | UCCCCGGCGC | CGCUCGUGGG | GGGCCCAAGU |
| 201 | CCUUCUGAUC | GAGGCCCAGC | CCGUGGACGG | UGUGAGGCCG | GUAGCGGCCC | CCGGCGCGCC | GGGCCCGGGU | CUUCCCGGAG | UCGGGUUGCU | UGGGAAUGCA |
| 301 | GCCCAAAGCG | GGUGGUAAAC | UCCAUCUAAG | GCUAAAUACC | GGCACGAGAC | CGAUAGUCAA | CAAGUACCGU | AAGGGAAAGU | UGAAAAGAAC | UUUGAAGAmGAm |
| 401 | GAGUUCAAGA | GGGCGUGAAA | CCGUUAAGAG | GUAAACGGGU | GGGGUCCGCG | CAGUCCGCCC | GGAGGAUUCA | ACCCGGCGGC | GGGUCCGGCC | GUGUCGGCGG |
| 501 | CCCGGCGGAU | CUUUCCCGCC | CCCCGUUCCU | CCCGACCCCU | CCACCCGCCC | UCCCUUCCCC | CGCCGCCCCU | CCUCCUCCUC | CCCGGAGGGG | GCGGGCUCCG |
| 601 | GCGGGUGCGG | GGGUGGGCGG | GCGGGGCCGG | GGGUGGGGUC | GGCGGGGGAC | CGUCCCCCGA | CCGGCGACCG | GCCGCCGCCG | GGCGCAUUUC | CACCGCGGCG |
| 701 | GUGCGCCGCG | ACCGGCUCCG | GGACGGCUGG | GAAGGCCCGG | CGGGGAAGGU | GGCUCGGGGG | GCCCCGUCCG | UCCGUCCGUC | CGUCCUCCUC | CUCCCCCGUC |
| 801 | UCCGCCCCCC | GGCCCCGCGU | CCUCCCUCGG | GAGGGCGCGC | GGGUCGGGGC | GGCGGCGGCG | GCGGCGGUGG | CGGCGGCGGC | GGCGGCGGCG | GGACCGAAAC |
| 901 | CCCCCCCGAG | UGUUACAGCC | CCCCCGGCAG | CAGCACUCGC | CGAAUCCCGG | GGCCGAGGGA | GCGAGACCCG | UCGCCGCGCU | CUCCCCCCUC | CCGGCGCCCA |
| 1001 | CCCCCGCGGG | GAAUCCCCCG | CGAGGGGGGU | CUCCCCCGCG | GGGGCGCGCC | GGCGUCUCCU | CGUGGGGGGG | CCGGGCCACC | CCUCCCACGG | CGCGACCGCU |
| 1101 | CUCCCACCCC | UCCUCCCCGC | GCCCCCGCCC | CGGCGACGGG | GGGGGUGCCG | CGCGCGGGUC | GGGGGGCGGG | GCGGACUGUC | CCCAGUGCGC | CCCGGGCGGG |
| 1201 | UCGCGCCGUC | GGGCCCGGGG | GAGGUUCUCU | CGGGGCCACG | CGCGCGUCCC | CCGAAGAGGG | GGACGGCGGA | GCGAGCGCAC | GGGGUCGGCG | GCGACGUCGG |
| 1301 | CUACCCACCC | GACCCGmUCUU | GAAmACAmCGGA | CCAAGGAGUCm | UAACACGUGC | GCGAGUCGGG | GGCUCGCACG | AAAGCCGCCG | UGGCGCAAUG | AAGGUGAAGG |
| 1401 | CCGGCGCGCU | CGCCGGCCGA | GGUGGGAUCC | CGAGGCCUCU | CCAGUCCGCC | GAGGGCGCAC | CACCGGCCCG | UCUCGCCCGC | CGCGCCGGGG | AGGUGGAGCA |
| 1501 | CGAGCGCACG | UGUUAGGACC | CGmAAmAGAUGG | UGAAmCΨAUGC | CUGGGCAGGG | CGAAGCCAGA | GGAAACUCUG | GUGGAGGUCC | GΨAGCGGUCC | UGACGUGCAA |
| 1601 | AUCGGUCGUC | CGACCUGGGU | AUAGGmGGCGA | AAGACUAAUC | GAACCAUCUA | GUAGCUGGUU | CCCUCCGAAG | UUUCCCΨCAG | GAΨAGCUGGC | GCUCUCGCAG |
| 1701 | ACCCGACGCA | CCCCCGCCAC | GCAGUUUUAU | CCGGUAAAGC | GAAΨGAUUAG | AGGUCUUGGGm | GCCGAAACGA | UCUmCAACCΨA | ΨΨCUCAAACU | UΨAAAUGGGU |
| 1801 | AAGAAGCCCG | GCUCGCUGGC | GUGGAGCCGG | GCGUGGAAUG | CGAGUGCCUA | GUGGGCCACΨ | UΨUGGUAAGC | AmGAACUGGCG | CmUGCGGGAUG. | AACCGAACGC |
| 1901 | CGGGUUAAGG | CGCCCGAUGC | CGACGCUCAU | CAGACCCCAG | AAAAGGUGUU | GGUUGAUAUA | GACAGCAGGA | CGGUGGCCAU | GGAAGUCGGA | AUCCGCUAAG |

2001 GAGUGUGUAA CAACUCACCU GCCGAAUCAA CUAGCCCUGA AAAUGGAUGG CGCUGGAGCG UCGGGCCCAU ACCCGGCCGU CGCCGGCAGU CGAGAGUGGA

2101 CGGGAGCGGC GGCGGCGGCG CGCGCGCGCG CGCGUGUGGU GUGCGUCGGA GGGCGGCGGC GGCGGCGGCG GCGGGGGUGU GGGGUCCUUC CCCCGCCCCC

2201 CCCCCCACGC CUCCUCCCCU CCUCCCGCCC ACGCCCCGCU CCCCGCCCCC GGAGCCCCGC GGACGCUACG CCGCGACGAG UAGGAGGGCC GCUGCGGUGA

2301 GCCUUGAAGC CUAGGGCGCG GGCCCGGGUG GAGCCGCCGC AGGUGCAGAU CmUUGGUGGUA GUAmGmCmAAAUA UUCAAACGAG AACUUUGAAG GCCGAAGUGG

2401 AmGAAGGGUUC CAUGUmGAACA GCmAGmUUGAAC AUGGGUCAGU CGGUCCUGAG AGAUGGGCGA GCGCCGUUCC GAAGGGACGG GCGAUGGCCU CCGUUGCCCU

2501 CGGCCGAΨCG AAAGGGAGUC GGGUUCAGAU CCCCGAAUCC GGAGUGGCGG AGAUGGGCGC CGCGAGGCGU CCAGUGCGGU AACGCGACCG AUCCCGGAGA

2601 AGCCGGCGGG AGCCCCGGGG AGAGUUCUCU UΨUCUUUGUG AAGGGCAGGG CGCCCUGGAA UGGGUUCGCC CCGAGAGAGG GGCCCGUGCC UUGGAAAGCG

2701 UCGCGGUUCC GGCGGCGUCC GGUGAGCUCU CGCUGGCCCU UGAAAAUCCG GGGGAGAGGG UGUAAAUCUC GCGCCGGGCC GUACCCAmUAU CCGCAGCAGG

2801 UCUCmCAAGGU GAACAmGCCUC UGGCmAUGUUG GAACAAUmGΨA GGΨAAGGGAA GUCGGCAAGC CmGGAUCCGUA ACUUCGmGGAU AAGGAUUGGC UCUAAGGGCU

2901 GGGUCGGUCG GGCUGGGGCG CGAAGCGGGG CUGGGCGCGC GCCGCGGCUG GACGAGGCGC CGCCGCCCCC CCCACGCCCG GGGCACCCCC CUCGCGGCCC

3001 UCCCCCGCCC CACCCCGCGC GCGCCGCUCG CUCCCUCCCC GCCCCGCGCC CUCUCUCUCU CUCUCUCCCC CGCUCCCCGU CCUCCCCCCU CCCCGGGGGA

3101 GCGCCGCGUG GGGGCGGCGG CGGGGGGAGA AGGGUCGGGG CGGCAGGGGC CGGCGGCGGC CCGCCGCGGG GCCCCGGCGG CGGGGGCACG GUCCCCCGCG

3201 AGGGGGGCCC GGGCACCCGG GGGGCCGGCG GCGGCGGCGA CUCUGGACGC GAGCCGGGCC CUUCCCGUGG AUCGCCCCAG CUGCGGCGGG CGUCGCGGCC

3301 GCCCCCGGGG AGCCCGGCGG GCGCCGGCGC GCCCCCCCCC CCACCCCACG UCUCGUCGCG CGCGCGUCCG CUGGGGGCGG GGAGCGGUCG GGCGGCGGCG

3401 GUCGGCGGGC GGCGGGGCGG GGCGGUUCGU CCCCCCGCCC UACCCCCCCG GCCCCGUCCG CCCCCCGUUC CCCCCUCCUC CUCGGCGCGC GGCGGCGGCG

3501 GCGGCAGGCG GCGGAGGGGC CGCGGGCCGG UCCCCCCCGC CGGGUCCGCC CCCGGGGCCG CGGUUCCGCG CGGCGCCUCG CCUCGGCCGG CGCCUAGCAG

3601 CCGACUUAGA ACUGGUGCGG ACCAGGGmGAA UCCGACΨGΨU UAAUUAAAAC AAAGCAUCGC GAAGGCCCGC GGCGGGUGUU GACGCGAUGU GAUUΨCUGCC

3701 CmAGUGCUCUG AAUGΨCAAmAG UGAAmGAAAUΨ CAAΨGAAGCG CGGGmUAAACG GCGGGAGΨAAm CΨAΨGACΨCΨ CUUAAGGUAG CCAAAmUGCCU CGmUCAUCUAA

3801 UUAGUGACmGC GCAUGAAΨmGG AΨGAAmCGAGAm UUCCCACUGU CmCCΨACCUAC ΨAΨCCAGCGA AACCACAmGCmC AAGGGAACGG GCUΨGGCmGGA AUCAGCGGGmG

3901 AAAGAAGACC CUGUUGAGCΨ UGACUmCUAGU CUGGCACGGU GAAGmAGACAU GAGAGGUGΨA GAAUAAGUGG GAGGCCCCCG GCGCCCCCCC GGUGUCCCCG

4001 CGAGGGGCCC GGGGCGGGGU CCGCCGGCCC UGCGGGCCGC CGmGUGAAAUA CCACmUACUCU GAUCGUUUUU UCACUGACCC GGUGAGGCGG GGGGGCGAGC

4101 CCCGAGGGGC UCUCGCUUCU GGCGCCAAGC GCCCGGCCGC GCGCCGGCCG GGCGCGACCC GCUCCGGGGA CAGUGCCAGG UGGGGAGUUU GACUGGmGGCG

4201 GUACACCUGU CAAACGGUAA CGCAGGUmGmUC CUAAGGCGAG CUCAGGGAGG ACAGAAACCU CCCGUGGAGC AGAAGGGCAA AAGCUCGCUU GAΨCUΨGAΨU

4301 UUCAGUmACGA AΨACAGACCG UGAAAGCGGG GCCUCACGAU CCUUCUGACC UUΨUGGGUUU ΨAAGCAGGAGm GUGUCAGAAA AGUUACCACA GGmGAUAACUG

4401 GCΨUGUGGCG GCCAAGCGUΨ CAΨAGCGACG ΨCGCUUUUUG AΨCCUUCGAU GUCGGCmΨCUU CCUAUCAUUG ΨGAAGCAGAA UUCACCAAGC GUΨGmGAUUmGmΨ

4501 UCACCCACUA AUAGGGAACG ΨGAmGCUGGGU UΨAGACmCGUC GUGAGACAGG UΨAGUUUUAC CCUACUGAUG AmUGUGΨUGΨU GCCAUGGUAAm UCCUGCUCAG

4601 UACGAGAGGA ACCGCAGGmUUm CAGmACAUΨUG GUGUAΨGmUGC. UUGGCUGAGG AGCCAAUGGG GCGAAGCUAC CAΨCUGUGGG AUUAUGACΨG AACGCCUCUA

4701 AGUCAGAAUC CCGCCCAGGC GGAACGAUAC GGCAGCGCCG CGGAGCCUCG GUUGGCCUCG GAUAGCCGGU CCCCCGCCUG UCCCCGCCGG CGGGCCGCCC

4801 CCCCCCUCCA CGCGCCCCGC GCGCGCGGGA GGGCGCGUGC CCCGCCGCGC GCCGGGACCG GGGUCCGGUG CGGAGUGCCC UUCGUCCUGG GAAACGGGGC

4901 GCGGCCGGAA AGGCGGCCGC CCCCUCGCCC GUCACGCACC GCACGUUCGU GGGGAACCUG GCGCUAAACC AΨΨCGUAGAC GACCUGCUUC UGGGUCGGGG

5001 ΨUUCGUACGΨ AGCAGAGCAG CUCCCUCGCU GCGAUCUAU UGAAAGUCAG CCCUCGACAC AAGGGUUUGU C

#### 5.8S rRNA sequencing with annotations of chemically modified nucleotides

1 CGACUCUUAG CGGUmGGAUCA CUCGGCUCGU GCGUCGAUGA AGAACGCAGC UAGCΨGCGAG AAUUAAUGΨG AAUUGmCAGGA CACAUUGAUC AUCGACACUU

101 CGAACGCACU UGCGGCCCCG GGUUCCUCCC GGGGCUACGC CUGUCUGAGC GUCGCUU

#### 5S rRNA sequencing (contains no chemical modifications)

1 GUCUACGGCC AUACCACCCU GAACGCGCCC GAUCUCGUCU GAUCUCGGAA GCUAAGCAGG GUCGGGCCUG GUUAGUACUU GGAUGGGAGA CCGCCUGGGA

101 AUACCGGGUG CUGUAGGCUU

**18S, 28S, 5.8S & 5S rRNA sequencing with annotations of chemically modified nucleotides**

A4910 is indicated in red (see Extended Data Fig. 12); sequences in grey were not covered.

**Extended Data Fig. 13**

| | | | |
|---|---|---|---|
| 18S_1F | CCTGGTTGATCCTGCCAGTAGC- | 18S_160R | CCACAGTTATCCAAGTAGG |
| 18S_544_F | TAACAATACAGGACTCTTTCG | 18S-972_R | TCCAAGAATTTCACCTCTAGC |
| 18S_1600_F | TAACCCGTTGAACCCCA TTCGTGAT | 18S-1581_R | GTTACCCGCGCCTGCCGGCG |
| | | 18S_1760_R | CCGCCCCCCGGCGGGGCCG |
| | | 18S_1860_R | TAATGATCCTTCCGCAGGTTCACC |

| | | | |
|---|---|---|---|
| 28S_1_F | 5'CGCGACCTCAGATCAGACGTGG3' | 28S-450-R | 5'ACTGCGCGGACCCCACCCGTTT3' |
| 28S_510_F | ATCTTTCCCGCCCCCCGTTCCT | 28S_1300_R | TTTCAAGACGGGTCGGGTGGGTAGCCGACGT |
| 28S_1500_F | GAACTATGCCTGGGCAGGGC | 28S-2260-R | TCCTACTCGTCGCGGCGTAGCGTCCGCGGGGCT |
| 28S_2030_F | AATCAACTAGCCCTGAAAATGGATG | 28S_2750_R | CGGCGCGAGATTTACACCCTCTC |
| 28S-2900-F | ATAAGGATTGGCTCTAAGGGCTGGGTCG GT | 28S-3270-R | ATCCACGGGAAGGGCCCGGCTCGCGTCCAGA |
| 28S-3620_F | GACCAGGGGAATCCGACTGTTTAATT | 28S_3600_R | CACCAGTTCTAAGTCGGCTGCTAG |
| 28S-4440_F | AAGCGTTCATAGCGACGTCGCTTTTTGAT | 28S_5060_R | GACAAACCCTTGTGTCGAGGGC |
| 28S_2750_F | GAGAGGGTGTAAATCTCGCGCCG | 28S_5000_R | GCTACGTACGAAACCCCGACC |
| | | 28S_4690_R | ATAATCCCACAGATGGTAGCTTCGCCCCATT |

| | | | |
|---|---|---|---|
| 5.8S_1_F | CGACTCTTAGCGGTGGATCACTC | 5.8S_157_R | AAGCGACGCTCAGACAGGC |

| | | | |
|---|---|---|---|
| 5S_1_F | GTCTACGGCCATACCACCCTG | 5S_120_R | AAGCCTACAGCACCCGGTATTC |

**Primer sequences used for human 18S, 28S, 5.8S and 5S rRNA sequencing**

**Extended Data Fig. 14**
